# Ventral Tegmental Area GABA, glutamate, and glutamate-GABA neurons are heterogenous in their electrophysiological and pharmacological properties

**DOI:** 10.1101/2020.07.22.216093

**Authors:** Jorge Miranda-Barrientos, Ian Chambers, Smriti Mongia, Bing Liu, Hui-Ling Wang, Gabriel E Mateo-Semidey, Elyssa B Margolis, Shiliang Zhang, Marisela Morales

## Abstract

The ventral tegmental area (VTA) contains dopamine neurons intermixed with GABA-releasing (expressing vesicular GABA transporter, VGaT), glutamate-releasing (expressing vesicular glutamate transporter, VGluT2), and co-releasing (co-expressing VGaT and VGluT2) neurons. By delivering INTRSECT viral vectors into VTA of double *vglut2-Cre/vgat-Flp* transgenic mice, we targeted specific VTA cell populations for ex vivo recordings. We found that VGluT2^+^ VGaT^−^ and VGluT2^+^ VGaT^+^ neurons on average had relatively hyperpolarized resting membrane voltage, greater rheobase, and lower spontaneous firing frequency compared to VGluT2^−^ VGaT^+^ neurons, suggesting that VTA glutamate-releasing and glutamate-GABA co-releasing neurons require stronger excitatory drive to fire than GABA-releasing neurons. In addition, we detected expression of Oprm1mRNA (encoding μ opioid receptors, MOR) in VGluT2^+^ VGaT^−^ and VGluT2^−^ VGaT^+^ neurons, and their hyperpolarization by the MOR agonist DAMGO. Collectively, we demonstrate the utility of the double transgenic mouse to access VTA glutamate, glutamate-GABA and GABA neurons, and show some electrophysiological heterogeneity among them.

**Impact Statement:** Some physiological properties of VTA glutamate-releasing and glutamate-GABA co-releasing neurons are distinct from those of VTA GABA-releasing neurons. μ-opioid receptor activation hyperpolarizes some VTA glutamate-releasing and some GABA-releasing neurons.

## Introduction

The ventral tegmental area (VTA) is a midbrain structure containing dopamine neurons that play a major role in motivated behaviors (Wise, 2004, Berridge et al., 2007, Bromberg-Martin et al., 2010). While it has classically been thought of as a dopaminergic structure, the VTA contains multiple types of neurons, including neurons that release GABA (expressing the synthesis enzyme glutamate decarboxylase, GAD, and the vesicular GABA transporter, VGaT) and neurons that release glutamate (expressing vesicular glutamate transporter type 2, VGluT2) (Yamaguchi et al., 2007). Moreover, we recently demonstrated that some VTA neurons release both glutamate and GABA (Root et al., 2014). These neurons express both VGluT2 and VGaT mRNA (VGluT2^+^ VGaT^+^ neurons), whose proteins are distributed in separate pools of synaptic vesicles within a common axon terminal (Root et al., 2018). We have found a lateromedial gradient of distribution for VGluT2^+^ VGaT^+^ neurons and those that release glutamate without GABA (VGluT2^+^ VGaT^−^ neurons) or release GABA without glutamate (VGluT2^−^ VGaT^+^ neurons) (Root et al., 2018).

The *ex vivo* electrophysiological properties of putative VTA dopamine neurons had been investigated for decades; the dopaminergic identification of VTA recorded neurons has been achieved by the immunocytochemical detection of tyrosine hydroxylase (TH) in rat (Margolis et al., 2006) or by *in vivo* labeling in transgenic mice (Khaliq & Bean, 2010). Characterization of VTA GABA neurons in *ex vivo* recordings has been achieved by GAD protein and mRNA detection in recorded neurons in rat (Margolis et al., 2012) or by *in vivo* labeling in transgenic mice with constitutive or viral vector-induced expression of green fluorescent protein (GFP) under the control of the GAD 2 (Tan et al., 2012) or VGaT (va Zessen et al., 2012) promoters. *Ex vivo* recordings of VTA glutamate neurons have been made in mice that constitutively express GFP under the control of the VGluT2 promoter (Hnasko et al., 2012). Importantly, these studies did not differentiate neurons that co-release glutamate and GABA (VGluT2^+^ VGaT^+^) from those that are only glutamate-releasing (VGluT2^+^ VGaT^−^) or GABA-releasing (VGluT2^−^ VGaT^+^), raising the possibility that some of the overlapping physiological properties reported in these groups specifically belong to this co-releasing population.

While existing transgenic mice allow *in vivo* tagging of the entire population of VTA neurons expressing VGluT2 or VGaT, these transgenic lines are not suitable for the selective tagging of VGluT2^+^ VGaT^+^, VGluT2^+^ VGaT^−^, or VGluT2^−^ VGaT^+^ neurons. To overcome this limitation, we applied an intersectional approach to induce expression of enhanced YFP (eYFP) in the different classes of VTA neurons (Fenno et al., 2014). We found that some electrophysiological properties do vary across these different classes of VTA neurons, and determined that not only VGluT2^−^ VGaT^+^ neurons, but also VGluT2^+^ VGaT^−^ neurons, were postsynaptically inhibited by MOR activation.

## Results

### Selective targeting of VTA VGluT2^+^ VGaT^+^, VGluT2^+^ VGaT^−^ and VGluT2^−^VGaT^+^ neurons

To tag VGluT2^+^ VGaT^+^, VGluT2^+^ VGaT^−^, and VGluT2^−^ VGaT^+^ neurons, we generated *vglut2-Cre/vgat-Flp* mice by crossing *vglut2-Cre* mice with *vgat-Flp* mice (Figure 1A) and injected INTRSECT adeno associated viral (AAV) vectors into their VTA (Fenno et al., 2014). In different cohorts of *vglut2-Cre/vgat-Flp* mice, we injected AAV-C_ON_/F_ON_-eYFP vectors (requiring Cre and Flp recombinases for eYFP expression) to target VGluT2^+^ VGaT^+^ neurons; AAV-C_ON_/F_OFF_-eYFP vectors (requiring the presence of Cre recombinase and absence of Flp recombinase for eYFP expression) to target VGluT2^+^ VGaT^−^ neurons; or AAV-C_OFF_/F_ON_-eYFP vectors (requiring the absence of Cre recombinase and presence of Flp recombinase for eYFP expression) to target VGluT2^−^ VGaT^+^ neurons (Figure 1B-C).

**Figure 1.**
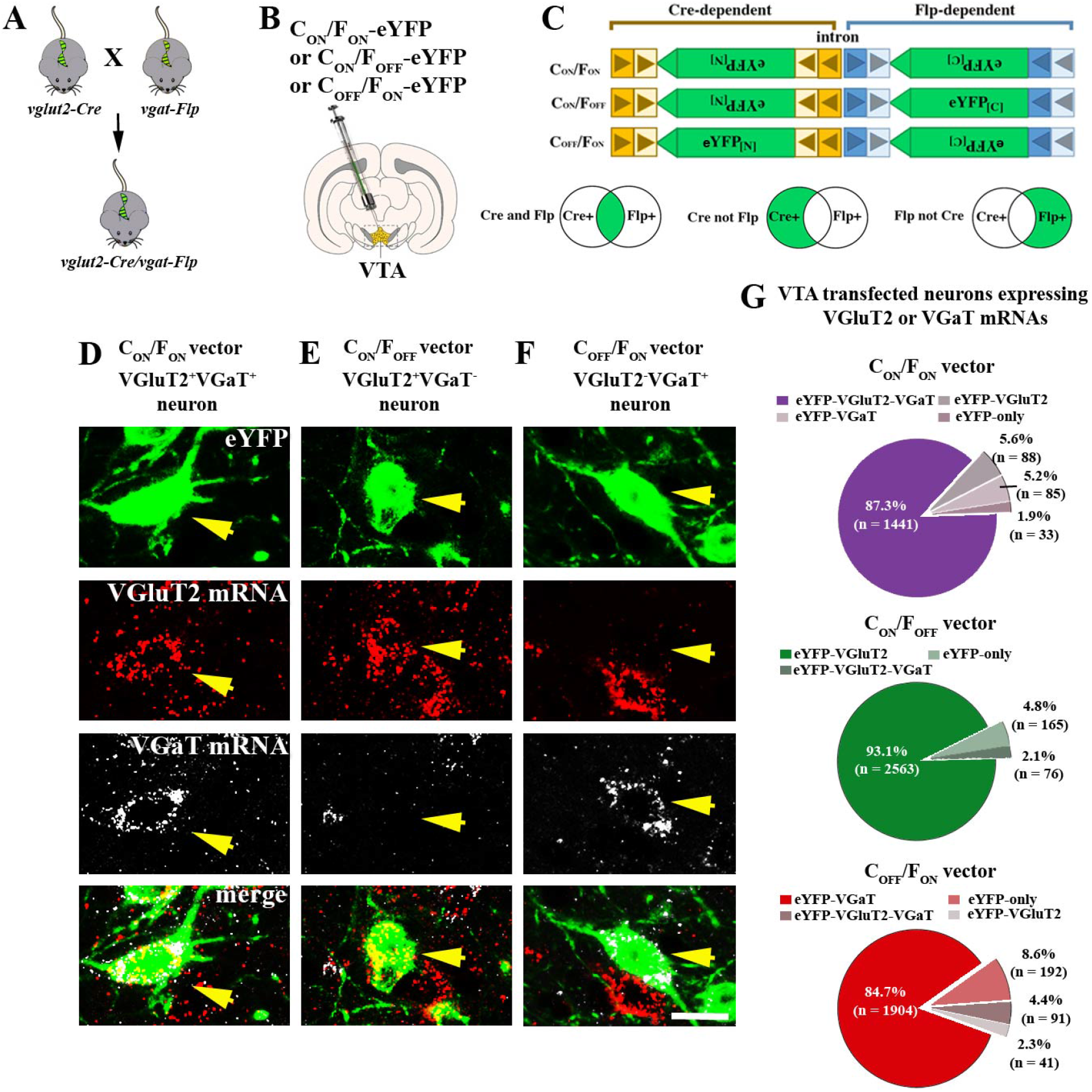
Selective targeting of VTA VGluT2^+^ VGaT^+^, VGluT2^+^ VGaT^−^ and VGluT2^−^VGaT^+^ neurons. **(A-C)** Crossing of *vglut2-Cre* and*vgat-Flp* mice to generate *vglut2-Cre/vgat-Flp* mouse and intra-VTA injections of INTRSECT AAV-CON/FON-eYFP to target VGluT2^+^ VGaT^+^ neurons, AAV-CON/FOFF-eYFP to target VGluT2^+^ VGaT^−^ neurons or AAV-COFF/FON-eYFP vectors to target VGluT2^−^ VGaT^+^ neurons. **(D)** Co-expression of VGluT2 mRNA and VGaT mRNA in VTA VGluT2^+^ VGaT^+^ eYFP neuron. **(E)** VGluT2 mRNA without VGaT mRNA in VTA VGluT2^+^ VGaT-eYFP neuron. **(F)** VGaT mRNA without VGluT2 mRNA in VTA VGluT2^−^VGaT^+^ eYFP neuron. Scale bar =20 uM **(G)** Total percentage of VTA transfected neurons co-expressing VGluT2 mRNA or VGaT mRNA (n= 3 mice per group).

After confirming VTA neuronal expression of eYFP in each cohort of mice (Figure 1D-F), we examined the mRNA expression of VGluT2 or VGaT within eYFP expressing neurons (Figure 1D-F). In the VTA of mice locally injected with AAV-C_ON_/F_ON_-eYFP vectors (to tag VGluT2^+^ VGaT^+^ neurons) (Figure 1D), we found that within the total population of eYFP expressing neurons (1,647 neurons, 3 mice; Figure 1G), approximately 90% expressed both VGluT2 and VGaT mRNAs (87.3% ± 2.4%; 1,441/1,647); 6% expressed only VGluT2 mRNA (5.6% ± 1.6%; 88/1,647 neurons), close to 5% expressed only VGaT mRNA (5.2% ± 2.0 %; 85/1,647, Figure 1G), and rarely lacked both VGluT2 and VGaT mRNAs (1.93% ± 0.35%; 33/1647). In the VTA of mice locally injected with AAV-C_ON_/F_OFF_-eYFP vectors (to tag VGluT2^+^ VGAT^−^ neurons) (Figure 1E), we found that within the total population of eYFP neurons (2,804 neurons, 3 mice; Figure 1G) more than 90% expressed VGluT2 mRNA (93.1% ± 2.8%, 2,563/2,804), none expressed VGaT mRNA alone, they rarely expressed VGluT2 mRNA together with VGaT mRNA (2.1% ± 0.9%, 76/2,804), and a small number lacked both VGluT2 or VGaT mRNAs (4.8% ± 2.0%, 165/2,804). In the VTA of mice locally injected with AAV-C_OFF_/F_ON_-eYFP vectors (to tag VGluT2^−^ VGaT^+^ neurons) (Figure 1F), we found that within the total population of eYFP neurons, close to 85% expressed VGaT mRNA (84.7% ± 2.1%, 1,904/2,228 neurons, 3 mice) (Figure 1F, G), rarely expressed VGluT2 mRNA (2.3% ± 1.1%, 41/2,228 neurons) (Figure 1G), infrequently had VGaT mRNA together with VGluT2 mRNA (4.4% ± 1.4%; 91/2,228 neurons) (Figure 1G), and a small number lacked both VGluT2 and VGaT mRNAs (8.6% ± 0.5%, 192/2,228 neurons) (Figure 1G). Collectively, these findings indicate that using *vglut2-Cre/vgat-Flp* mice in combination with the tested INTRSECT viral vectors for the selective tagging of the three classes of VTA neurons generates very few false positive classifications.

### Intrinsic properties of VTA VGluT2^+^ VGaT^+^, VGluT2^+^ VGaT^−^ and VGluT2^−^VGaT^+^ neurons

Next we determined the spontaneous action potential (AP) activity of the three classes of neurons with cell attached recordings in horizontal brain slices. We detected spontaneous activity in 46.2% of the VGluT2^+^ VGaT^+^ neurons (24/52 neurons, 28 mice), 52.6% of the VGluT2^+^ VGaT^−^ neurons (24/49 neurons, 27 mice) and 70.7% of the VGluT2^−^ VGaT^+^ neurons (29/41 neurons, 19 mice) (Figure 2C). Within the spontaneously active neurons (77 neurons), we found that the mean firing frequency was lower in VGluT2^+^ VGaT^+^ and VGluT2^+^VGaT^−^ neurons than in VGluT2^−^ VGaT^+^ neurons (3.4 ± 0.7 Hz for VGluT2^+^ VGaT^+^, n= 24 neurons, 20 mice; 3.5 ± 0.7 Hz for VGluT2^+^ VGaT^−^, n= 24 neurons, 18 mice; and 8.3 ± 2.1 Hz for VGluT2^−^ VGaT^+^ n= 29 neurons, 14 mice) (Figure 2B, D). Together these findings indicate that more VGluT2^−^ VGaT^+^ neurons fire spontaneously, and on average they fire faster, than VGluT2^+^ VGaT^+^ or VGluT2^+^VGaT^−^ neurons. Next, we analyzed the extracellular AP durations from these neurons and found that VGluT2^+^ VGaT^+^ neurons have similar AP durations to VGluT2^−^ VGaT^+^ neurons, and both of these phenotypes have shorter AP durations than VGluT2^+^ VGaT^−^ neurons (1.3 ± 0.1 ms for VGluT2^+^ VGaT^+^, n= 24 neurons, 20 mice; 1.7 ± 0.1 ms for VGluT2^+^ VGaT^−^, n= 24 neurons, 18 mice; and 1.2 ± 0.1 ms for VGluT2^−^ VGaT^+^ n= 29 neurons, 14 mice) (Figure 2F,G). Because VTA neurons are known to have pacemaker activity as well as burst firing patterns, we also evaluated the regularity of firing. We analyzed the coefficient of variation (CV) of inter-spike intervals (ISIs), and found that both VGluT2^+^ VGaT^+^ and VGluT2^+^VGaT^−^ neurons had higher CVs of ISIs compared to VGluT2^−^ VGaT^+^ neurons (48.5 ± 6.00 for VGluT2^+^ VGaT^+^, 41.8 ± 6.5 for VGluT2^+^ VGaT^−^, and 37.2 ± 4.9 for VGluT2^−^ VGaT^+^) (Figure 2E). These findings indicate that the spontaneous firing of VGluT2^+^ VGaT^+^ and VGluT2^+^VGaT^−^ neurons is more irregular than that of VGluT2^−^ VGaT^+^ neurons.

**Figure 2.**
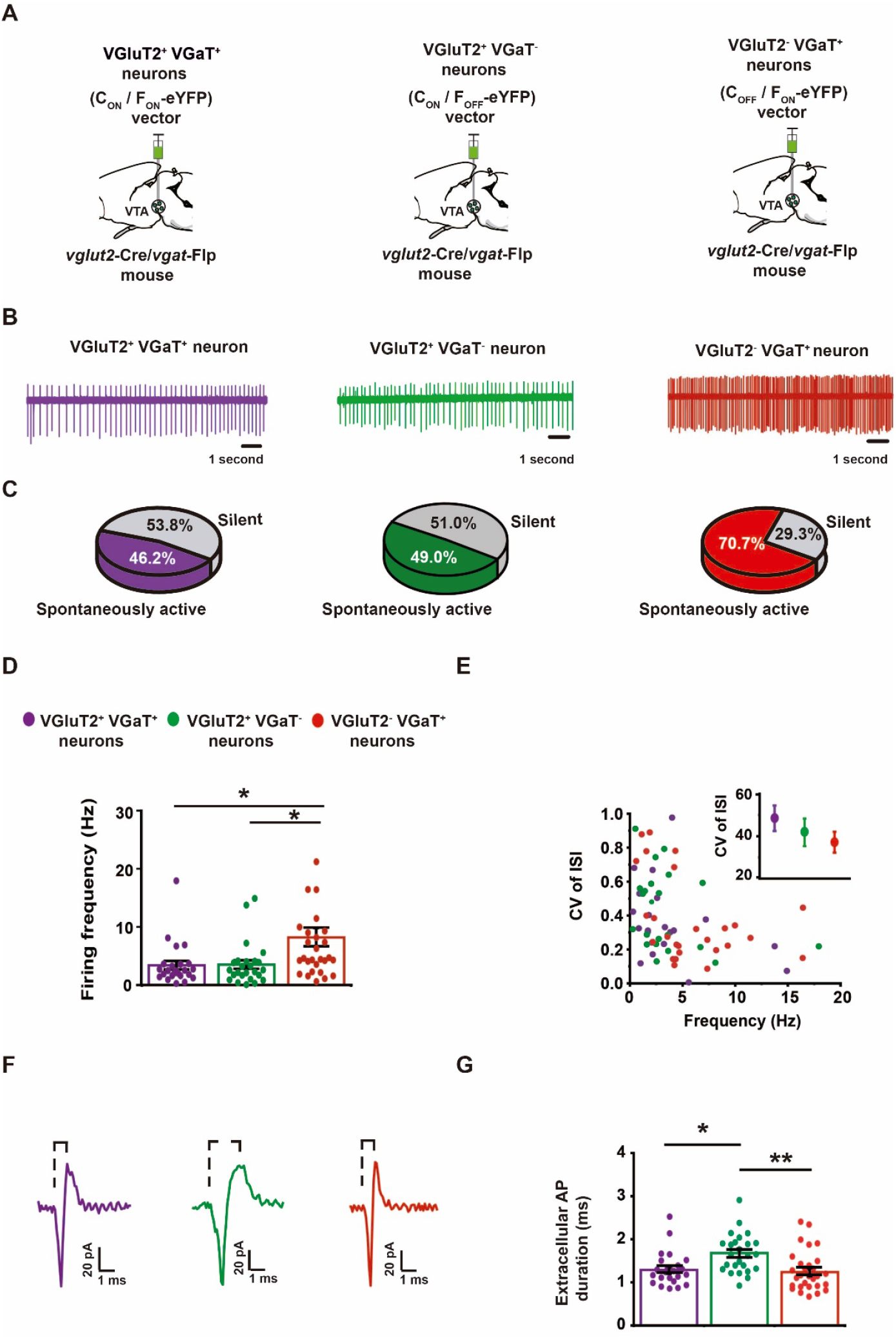
Spontaneous firing activity in VTA VGluT2^+^ VGaT^+^, VGluT2^+^ VGaT^−^, and VGluT2^−^VGaT^+^ neurons during *ex vivo* cell attached recordings. **(A)** Schematic representation of intra-VTA injections of vectors (AAV-CON/FON-eYFP, AAV-CON/FOFF-eYFP and AAV-COFF/FON-eYFP) in *vglut2-Cre*/*vgat-Flp* mice. **(B)** Traces recorded in the cell attached configuration in horizontal slices from identified spontaneously active VTA neurons. **(C)** Proportion of VTA spontaneously active vs quiescent neurons. **(D)** Summary of spontaneous firing rate across VTA neurons. VGluT2^−^VGaT^+^ neurons have higher firing frequencies than VGluT2^+^ VGaT^+^ and VGluT2^+^ VGaT^−^ neurons. One-way ANOVA F2,77 = 5.795 p= 0.046; Tukey’s *post hoc* test *p˂ 0.05. (E) There is no relationship between firing frequency and coefficient of variation (CV) of inter-spike intervals (ISIs) for these neuronal types. Inset, summary of CV of ISIs in VGluT2^+^ VGaT^+^, VGluT2^+^ VGaT^−^, and VGluT2^−^VGaT^+^ neurons. Example traces (F) and summary durations (G) of extracellular recorded APs. One-way ANOVA F2,77 = 6.745= 0.0026; Tukey’s *post hoc* test *p˂ 0.05, **p˂0.01

Next, using whole-cell recordings, we examined intrinsic electrophysiological properties of these three classes of VTA neurons. We found that the resting membrane potential was −68.0 ± 1.2 mV for VGluT2^+^ VGaT^+^ neurons (n= 52 neurons, 27 mice), −64.6 ± 0.9 mV for VGluT2^+^ VGaT^−^ neurons (n= 49 neurons, 27 mice), and −59.6 ± 0.9 mV for VGluT2^−^ VGaT^+^ neurons (n= 41 neurons, 19 mice). We detected the greatest rheobase in VGluT2^+^ VGaT^−^ neurons (37.3 ± 5.6 pA, n= 48 neurons, 27 mice), followed by VGluT2^+^ VGaT^+^ neurons (29.0 ± 5.5 pA, n= 52 neurons, 28 mice), and the lowest rheobase in VGluT2^−^ VGaT^+^ neurons (14.9 ± 2.9 pA, n= 41, 19 mice) (Table 1). Collectively these findings indicate that VTA neurons expressing VGluT2, with or without VGaT, are less excitable than VTA neurons lacking VGluT2, and the VGluT2^+^ VGaT^−^ neurons are the least excitable among the 3 classes. Across the three classes of VTA neurons, there were no differences in membrane capacitance, membrane resistance, membrane time constant, or AP threshold. However, in line with our cell attached data, we found that VGluT2^+^ VGaT^−^ neurons have longer duration APs than VGluT2^+^ VGaT^+^ and VGluT2^−^ VGaT^+^ neurons (2.2 ± 0.1 ms for VGluT2^+^ VGaT^+^, n= 52 neurons, mice; 2.9± 0.2 ms for VGluT2^+^ VGaT^−^, n= 48 neurons, 27 mice and 1.8 ± 0.1 ms for VGluT2^−^ VGaT^+^, n= 41 neurons, 19 mice) (Table 1).

**Table 1.** Intrinsic membrane and action potential properties in VTA VGluT2^+^ VGaT^+^, VGluT2^+^ VGaT^−^, and VGluT2^−^VGaT^+^ neurons. Action potential (AP). Difference from membrane potential and action potential threshold (Gap). Millivolts (mV). Megaohms (MOhms). Picofarads (pF). Picoamperes (pA). Milliseconds (ms). Membrane potential: One-way ANOVA F2,140 = 16.65 p˂ 0.0001; Tukey’s *pot hoc* test *VGluT2^+^ VGaT^+^ vs VGluT2^+^ VGaT-p˂ 0.05, **VGluT2^+^ VGaT-vs VGluT2^−^VGaT^+^ p˂ 0.01, ***VGluT2^+^ VGaT^+^ vs VGluT2^−^ VGaT^+^ p˂ 0.001. Rheobase: One-way ANOVA F2,140 = 4.571 p= 0.0120; Tukey’s *pot hoc* test **VGluT2^+^ VGaT^−^ vs VGluT2^−^ VGaT^+^ p˂ 0.01. Gap: One-way ANOVA F2,130 = 12.37 p˂ 0.0001; Tukey’s *pot hoc* test ***VGluT2^+^ VGaT^+^ vs VGluT2^−^VGaT^+^ p˂ 0.001, ***VGluT2^+^ VGaT-vs VGluT2^−^VGaT^+^ p˂ 0.001. AP duration: One-way ANOVA F2,140 = 17.26 p˂ 0.0001; Tukey’s *pot hoc* test **VGluT2^+^ VGaT^+^ vs VGluT2^+^ VGaT-p˂ 0.001, ***VGluT2^+^ VGaT-vs VGluT2^−^VGaT^+^ p˂ 0.00.

| Class of VTA neuron | *Membrane potential (mV) | Membrane resistance (MOhms) | Capacitance (pF) | *Rheobase (pA) | AP Threshold (mV) | Gap (mV) | AP amplitude (mV) | *AP duration (ms) |
| --- | --- | --- | --- | --- | --- | --- | --- | --- |
| VGluT2<sup>+</sup> VGaT<sup>+</sup> (N= 52) | -68.0 ± 1.2 | 644.7 ± 68.4 | 30.9 ± 2.0 | 29.0 ± 5.5 | -39.1 ± 0.5 | 28.8 ± 1.3 | 70.7 ± 1.3 | 2.2 ± 0.1 |
| VGluT2<sup>+</sup> VGaT<sup>-</sup> (N=48) | -64.6 ± 0.9 | 655.7 ± 42.5 | 26.6 ± 1.3 | 37.3 ± 5.6 | -36.7 ± 0.7 | 28.0 ± 1.0 | 69.7 ± 1.0 | 2.9 ± 0.2 |
| VGluT2<sup>-</sup> VGaT<sup>+</sup> (N=41) | -59.6 ± 0.9 | 549.7 ± 53.7 | 30.6 ± 1.6 | 14.9 ± 2.9 | -38.4 ± 0.8 | 21.1 ± 0.9 | 72.9 ± 1.6 | 1.8 ± 0.1 |

Given that hyperpolarization-activated cation currents (*I*_h_) are present in both dopamine and non-dopamine neurons (Jones and Kauer, 1999; Margolis et al., 2006; Margolis et al., 2012; Hnasko et al., 2012), we tested the three classes of VTA neurons for *I*_h_. We detected *I*_h_ in 46.2% of VGluT2^+^ VGaT^+^ neurons (24/52 neurons, 28 mice), 53.3% of VGluT2^+^ VGaT^−^ neurons (24/45 neurons from 24 mice) and 92% of VGluT2^−^ VGaT^+^ neurons (35/38 neurons, 18 mice) (Figure 3B). While there was a wide range of *I*_h_ magnitude among each cell type, the mean *I*_h_ magnitude was smaller in VGluT2^+^ VGaT^+^ neurons compared to VGluT2^+^ VGaT^−^ neurons or VGluT2^−^ VGaT^+^ neurons (22.4 ± 3.2 pA for VGluT2^+^ VGaT^+^, 56.8 ± 12.2 pA for VGluT2^+^ VGaT^−^, and 48.3 ± 8.7 pA for VGluT2^−^ VGaT^+^) (Figure 3A, C). We found that regardless of the neuronal cell type, the neurons with larger *I*_h_ magnitudes were located in the lateral VTA and those with low amplitudes were in the medial VTA. These findings support a VTA latero-medial neuronal topography among dopamine and non-dopamine neurons.

**Figure 3.**
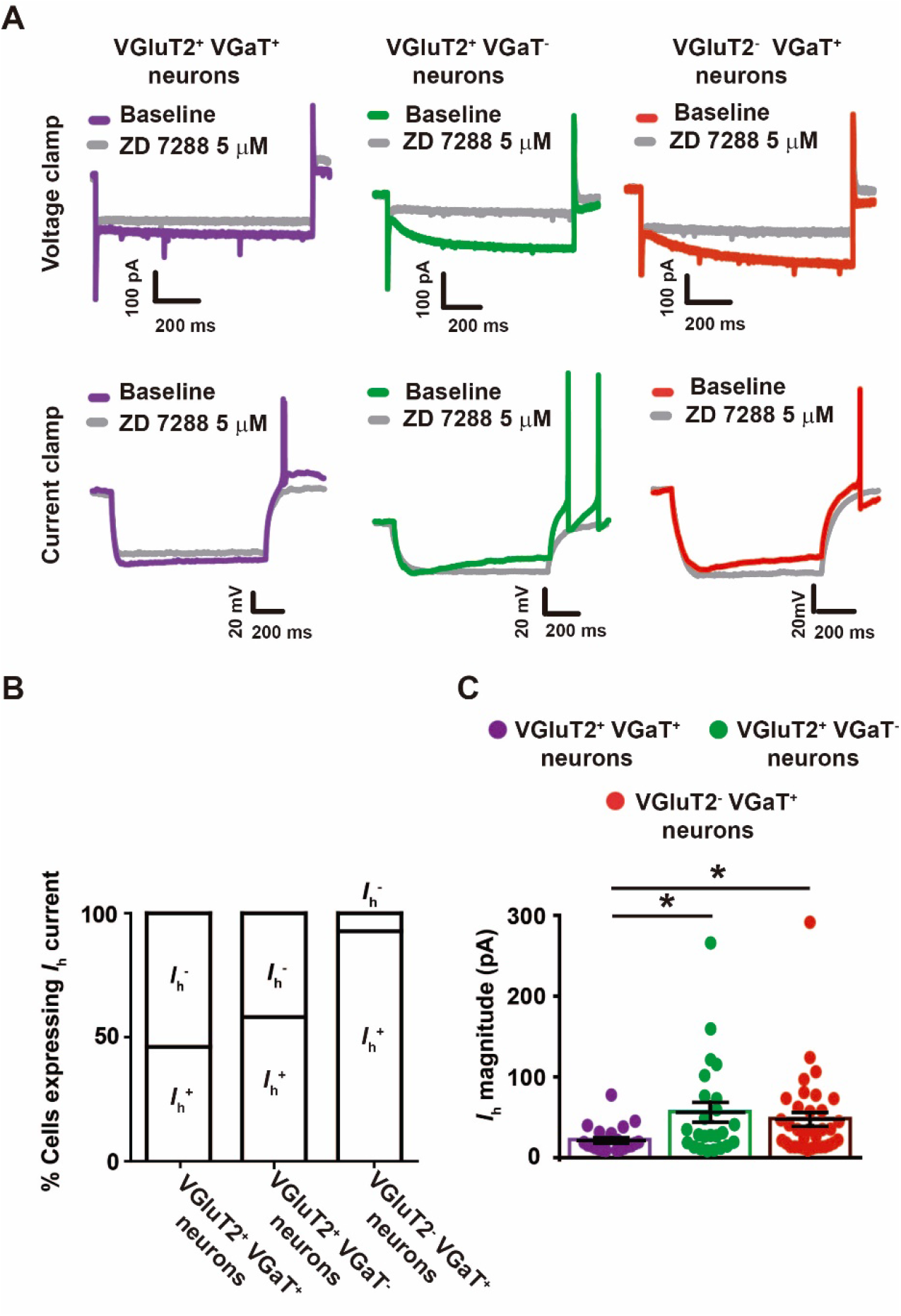
Many VTA VGluT2^+^ VGaT^+^, VGluT2^+^ VGaT^−^, and VGluT2^−^VGaT^+^ neurons have an *I*_**h**_. **(A)** Voltage clamp (top; step from −60 mV to −120 mV, 1s duration) and current clamp (bottom; step from 0 pA to 50-100 pA to reach mV= −120 mV) traces of *I*_h_ measurements in VTA neurons. ***I*_h_** was blocked by ZD 7288 (5 μM; gray traces). **(B)** Proportion of each VTA neuronal types that expressed *I*_h_. **(C)** Smaller *I* h amplitude was observed in VGluT2^+^ VGaT^+^ neurons compared to VGluT2^+^ VGaT-or VGluT2^−^VGaT^+^ neurons. One-way ANOVA F2,82= 3.528 p= 0.034, Tukey’s *post hoc* test *p˂ 0.05.

### Stimulated firing patterns of VTA VGluT2^+^ VGaT^+^, VGluT2^+^ VGaT^−^ and VGluT2^−^VGaT^+^ neurons

We next examined the stimulated firing patterns of these three classes of neurons by inducing AP firing in current clamp with a series of depolarizing current steps (500 ms, 10-150 pA).

We found that most VGluT2^+^ VGaT^+^ neurons (41/48 neurons, 24 mice) fired APs with short latency from initiation of the current pulse (159 ± 14 ms; Figure 4A-B, D). A small group of neurons responded to the injected current with delayed firing (7/48 neurons, 6 mice; 383 ± 47 ms; Figure 4A-B, D), displaying a slow depolarizing ramp prior to the first AP (Figure 4A). We found that the rheobase was generally greater in VGluT2^+^ VGaT^+^ neurons with long latency (101.4 ± 24.8 pA) than in those with short latency (17.8 ± 2.5 pA; Figure 4C). We observed that current injections below 100 pA produced depolarization block in some of the short latency neurons (14/41 neurons, 8 mice; Figure 4D). Across all short latency VGluT2^+^ VGaT^+^ neurons (41 neurons), more than half of them fired APs continuously during the entire current injection (27/41 neurons, 18 mice; Figure 4D). Furthermore, many of these neurons with continuous AP firing showed frequency adaptation (being more evident at currents above 100 pA; 18/27 neurons, 12 mice; Figure 4D), with the ISI increasing after each AP (Figure 4-figure supplement 1), but others either lacked or had minimal frequency adaptation (9/27 neurons, 7 mice; Figure 4-figure supplement 1) that permitted higher firing rates (38.7 ± 3.2 Hz sustained firing with adaptation; 84.2 ± 5.7 Hz for sustained firing without adaptation). VGluT2^+^ VGaT^−^ neurons were similar to VGluT2^+^ VGaT^+^ neurons, with short latency AP firing in response to injected current (29/45 neurons, 22 mice; 122 ± 16 ms latency) and fewer long latency responses with a depolarizing ramp leading to firing (16/45 neurons, 13 mice; 462 ± 11 ms latency; Figure 4A-B, D). Among the neurons with depolarizing ramp responses (VGluT2^+^ VGaT^+^ and VGluT2^+^ VGaT^−^ neurons), we detected A-type K^+^ currents (*I*_A_), which were blocked by the *I*_A_ blocker 4-Aminopyridine (4-AP; 2 mM) (Figure 4-figure supplement 2A-B). In addition, we found that 4-AP application decreased AP firing latency (4 VGluT2^+^ VGaT^+^ neurons and 4 VGluT2^+^ VGaT^−^ neurons, 8 mice; Baseline= 458 ± 7 ms, 4-AP= 192 ± 29 ms; Figure 4-figure supplement 2F), and increased the total number of APs fired during an input/output curve (10-150 pA, 500 ms) (4 VGluT2^+^ VGaT^+^ neurons and 4 VGluT2^+^ VGaT^−^ neurons, 8 mice; Baseline= 52.6 ± 17.8, 4-AP= 101 ± 24.3; Figure 4-figure supplement 2 C, G). Compared to any neurons expressing VGluT2 (VGaT^+^ or VGaT^−^), we found that all recorded VGluT2^−^ VGaT^+^ neurons had short latency AP firing responses to injected depolarizing current steps (38 neurons, 18 mice, 127.7 ± 13.5ms latency; Figure 4A-B, D), and a subset showed frequency adaptation (22/38 neurons, 15 mice) (Figure 4D). Collectively, these results demonstrate that both glutamate-GABA co-releasing (VGluT2^+^ VGaT^+^) and glutamate-releasing (VGluT2^+^ VGaT^−^) neurons are more heterogeneous in their firing properties than GABA-releasing (VGluT2^−^ VGaT^+^) neurons.

**Figure 4.**
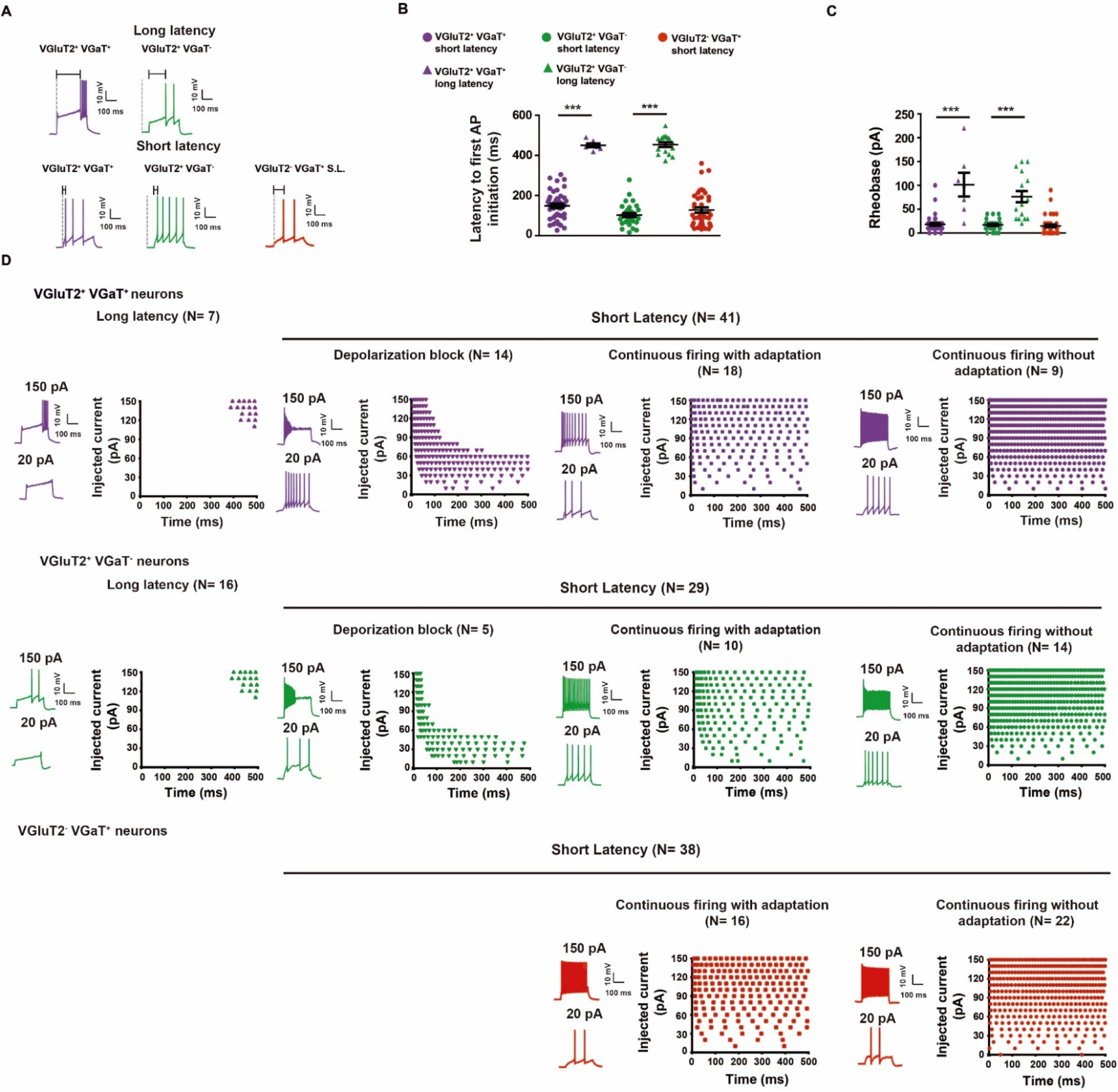
Stimulated firing patterns of VTA VGluT2^+^ VGaT^+^, VGluT2^+^ VGaT^−^, and VGluT2^−^VGaT^+^ neurons. **(A)** Example current clamp traces of VTA neurons with long or short latency AP firing in response to injected current steps. **(B)** VGluT2^+^ VGaT^+^, VGluT2^+^ VGaT^−^, and VGluT2^−^VGaT^+^ neurons with short or long latency to first AP. One-way ANOVA F4,139= 107.8 p˂0.0001, Tukey’s *post hoc* test ***p˂ 0.0001 **(C)**E voked AP firing in VGluT2^+^ VGaT^+^, VGluT2^+^ VGaT^−^, and VGluT2^−^VGaT^+^ neurons. Neurons with long latency to firing mostly required current injections ˃40 pA to drive firing. One-way ANOVA F4,13= 32.49 p˂0.0001, Tukey’s *post hoc* test ***p˂ 0.0001. **(D)** Example firing patterns in response to injected current steps sorted by long or short latency to firing. Short latency firing VGluT2^+^ VGaT^+^ and VGluT2^+^ VGaT-neurons go into depolarization block, show continuous firing with adaptation, or show continuous firing without adaptation. VGluT2^−^VGaT^+^ neurons only show responses of continuous firing with or without adaptation.

Given that previous studies have shown hyperpolarization-induced rebound burst firing in a subset of VTA dopamine and non-dopamine neurons mediated by *I*_h_ (Tateno and Robinson, 2011) or T-type calcium channels (Tracy et al., 2018, Woodward et al., 2019), we next tested the extent to which rebound firing occurs in the three classes of VTA neurons. After holding the resting membrane potential at −100 mV for 1 sec and then releasing that clamp, approximately 25% of VGluT2^+^ VGaT^+^ neurons showed rebound firing with short bursts of 2-4 APs (12/48 neurons, 10 mice; Figure 5), and this response was stable over repeated trials (Figure 5-suplement figure 1). In some of these neurons we tested if *I*_h_ blocker ZD 7288 stopped rebound firing, and found that it did in 2/4 neurons. In a different set of neurons we tested if mibefradil blocked the rebound firing, and found that it did in 3/5 neurons by the *I*_h_ blocker ZD 7288 (2/4 tested neurons) while in others it was blocked by T-type calcium channel blocker (Mibefradil; 3/5 tested neurons). Furthermore, we observed that 35% of VGluT2^+^ VGaT^−^ neurons (16/45 neurons, 11 mice) had rebound firing that was blocked by ZD 7288 (7/7 tested neurons) or Mibefradil (2/4 tested neurons) (Figure 5). In contrast, we found that half (20/38 neurons, 14 mice) of the VGluT2^−^ VGaT^+^ neurons showed rebound firing, which was blocked by ZD 7288 (5/6 tested neurons) or Mibefradil (2/5 tested neurons) (Figure 5). These findings demonstrated that some of the glutamate-GABA co-releasing neurons, glutamate-releasing and GABA-releasing neurons have rebound firing mediated by either *I*_h_ or T-type calcium channels.

**Figure 5.**
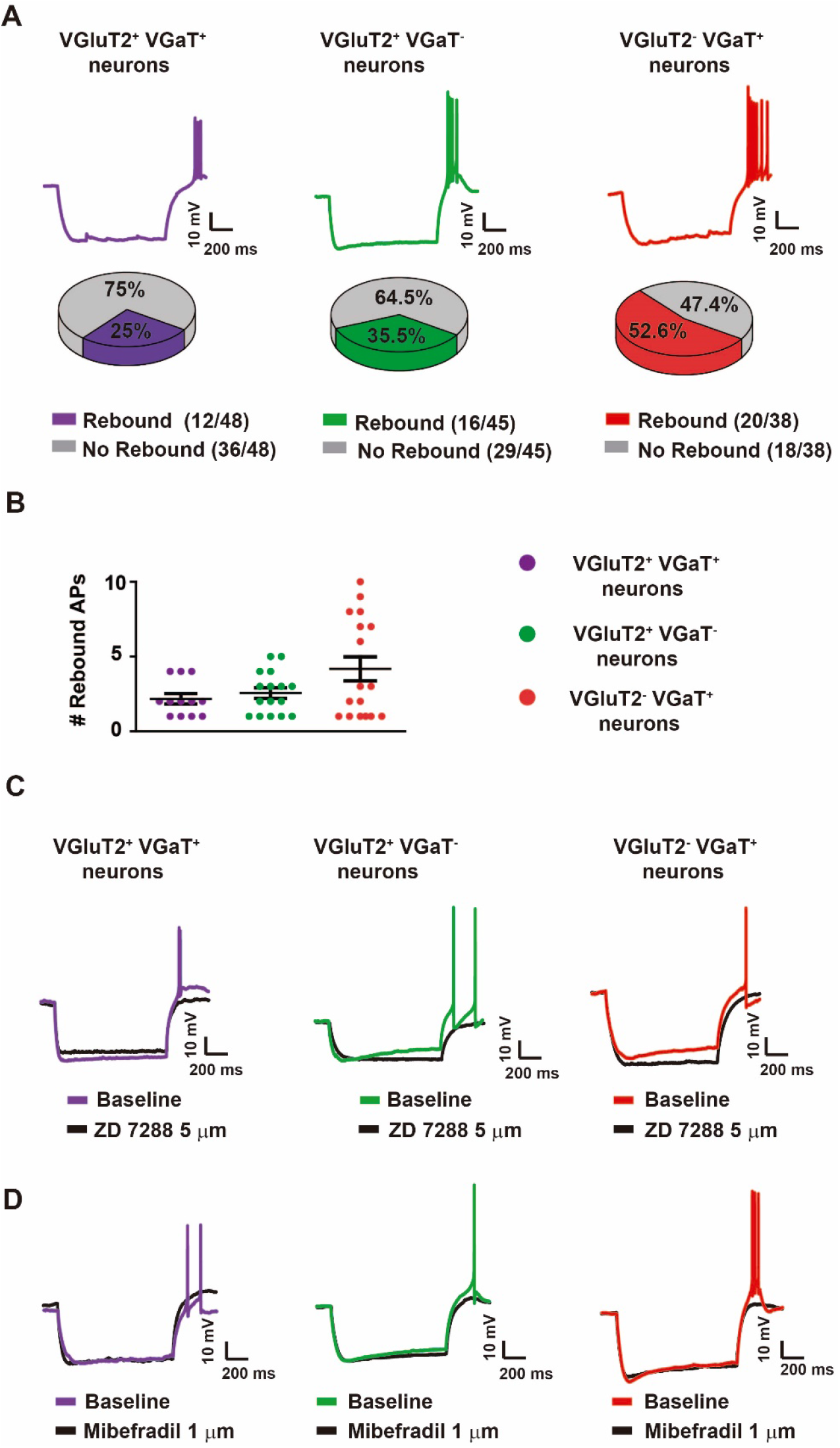
Rebound firing neurons is mediated by *I*_h_ and T-type calcium channels in VGluT2^+^ VGaT^+^, VGluT2^+^ VGaT^−^, and VGluT2^−^VGaT^+^ neurons. **(A)** Example current clamp traces (top) and proportion of neurons showing rebound firing (bottom) among VGluT2^+^ VGaT^+^ (purple), VGluT2^+^ VGaT^−^(green), and VGluT2^−^VGaT^+^ (red) neurons. **(B)** Number of rebound APs fired by VGluT2^+^ VGaT^+^ (purple), VGluT2^+^ VGaT^−^(green), and VGluT2^−^VGaT^+^ (red) neurons. **(C)** Example recordings in which rebound firing was blocked by ZD 7288 (5uM) (black traces). **(D)** Example recordings in which rebound firing was blocked by Mibefradil (1uM) (black traces).

### VTA neuronal clustering by electrophysiological properties

We next evaluated the extent to which the electrophysiological properties described above grouped together by applying a cluster analysis based on K-mean clustering and similarity parametric (Pearson’s correlation coefficient) on data obtained from the firing properties and intrinsic properties (Figure 6A). Based on the cluster analysis, we found 4 clusters of neurons with distinctive firing features, but these clusters did not uniquely correspond to any of our three neurotransmitter phenotypes (Figure 6B). For the 131 clustered neurons, 36.6 % (48/131 neurons, 33 mice) grouped in cluster 1, 32.1% (42/131 neurons, 31 mice) in cluster 2, 13.7% (18/131 neurons, 11 mice) in cluster 3, and 17.6% (23/131 neurons, 20 mice) in cluster 4 (Figure 6B).

**Figure 6.**
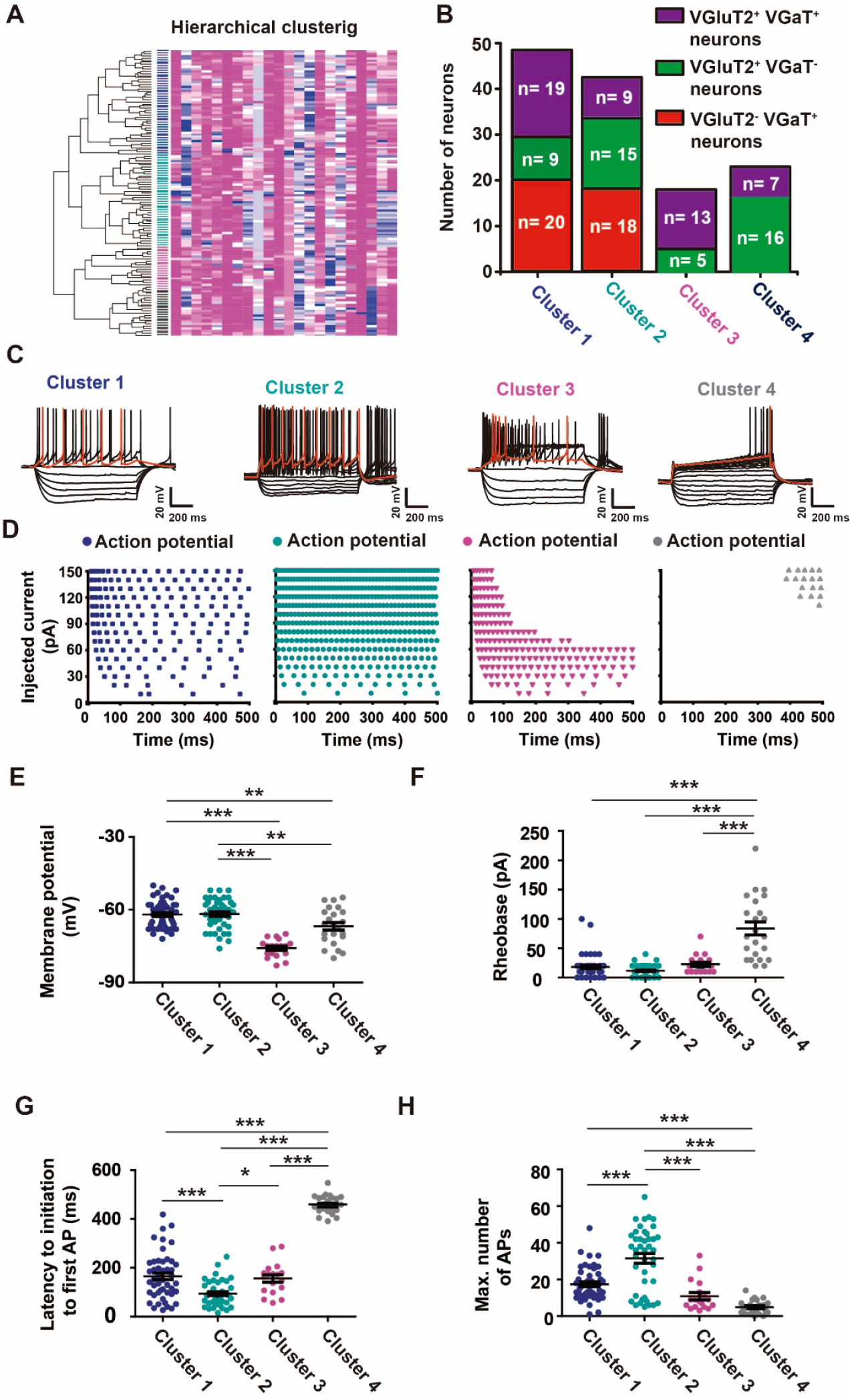
Hierarchical cluster analysis of electrophysiological properties of VTA VGluT2^+^ VGaT^+^, VGluT2^+^ VGaT^−^, and VGluT2^−^VGaT^+^ neurons. **(A)** Dendrogram and heat map of VTA neuronal electrophysiological properties. **(B)** Neurotransmitter neuronal phenotype distributions across clusters. **(C)** Example current clamp traces from neurons within each cluster in response to hyperpolarizing and depolarizing current injections. **(D)** Examples of firing patterns over time during depolarization steps in neurons within each cluster. **(E)** Membrane potentials of neurons in each cluster. One-way ANOVA F3,130= 28.25 p˂0.0001, Tukey’s *post hoc* test **p˂0.001 ***p˂ 0.0001. **(F)** R heobase of neurons in each cluster. One-way ANOVA F3,130= 42.45 p˂0.0001, Tukey’s *post hoc* test ***p˂ 0.0001. (G) Latency to fire APs in response to depolarizing current steps in neurons in each cluster. One-way ANOVA F3,130= 134.9 p˂0.0001, Tukey’s *post hoc* test *p˂ 0.05 ***p˂ 0.0001. **(H)** Maximum number of APs fired during depolarizing current steps (500 ms) of neurons in each cluster. One-way ANOVA F3,130= 30.64 p˂0.0001, Tukey’s *post hoc* test *p˂ 0.0001.

In cluster 1 (48 neurons), 39.5% were VGluT2^+^ VGaT^+^ (19/48 neurons, 12 mice), 18.8% were VGluT2^+^ VGaT^−^ (9/48 neurons, 7 mice), and 41.6% were VGluT2^−^ VGaT^+^ (20/48 neurons, 15 mice) (Figure 6B). Neurons grouped within the cluster 1 were characterized by a sustained AP firing with marked adaptation (46.5 ± 2.8 Hz maximal AP frequency, Figure 5H), and depolarized membrane potential (−61.9 ± 0.8 mV; Figure 6E). We found that 75% of the neurons within this cluster were spontaneously active during cell attached recordings (36/48 neurons, 29 mice), representing the highest percentage of spontaneously active neurons among the four different clusters. In cluster 2, 21.4% were VGluT2^+^ VGaT^+^ (9/42 neurons, 8 mice), 35.7% were VGluT2^+^ VGaT^−^ (15 /42 neurons, 12 mice), and 42.8% were VGluT2^−^ VGaT^+^ neurons (18/42 neurons, 12 mice) (Figure 6B). The neurons in this cluster had small rheobase (11.6 ± 1.4 pA; Figure 6F), short latency AP firing in response to depolarizing current steps (97 ± 10 ms Figure 6G), and sustained firing with minimal frequency adaptation in response to current steps (76.1 ± 3.4 Hz; Figure 6H). We found that 69% (29 /42 neurons, 23 mice) of the neurons in this cluster showed rebound firing after release from clamp induced hyperpolarization. In cluster 3, 72.2% were VGluT2^+^ VGaT^+^ (13/18 neurons, 8 mice) and 27.8% were VGluT2^+^ VGaT^−^ neurons (5/18 neurons, 4 mice; Figure 6B). Neurons grouped in this cluster exhibited rapid depolarization block during low magnitude depolarizing current steps (Figure 6C, D). In cluster 4, 30.4% (5/23 neurons from 5 mice) were VGluT2^+^ VGaT^+^ and 69.5% were VGluT2^+^ VGaT^−^ neurons (16 /23 neurons, 14 mice). No VGluT2^−^ VGaT^+^ neurons were classified to this cluster. Neurons in cluster 4 had larger rheobase (83.9 ± 11.1 pA; Figure 6F), long latencies to AP firing in response to injected current steps (452 ± 11 ms; Figure 6G), a depolarizing ramp before the onset of AP firing during current steps (Figure 6C-E), and included some neurons with *I*_A_ (Supplementary Figure 2C-G). We found that all neurons within cluster 4 were quiescent during cell attached recordings.

Next, we determined whether the 4 identified clusters of physiological properties had specific topography within the VTA by mapping the distribution of neurons that we filled with biocytin after recordings. We found cluster 1 neurons in ventral and middle VTA, with a dorsal to ventral, and lateral to medial increasing gradient of distribution (Figure 7B). We frequently observed cluster 2 neurons in middle VTA concentrated more medially (Figure 7C), cluster 3 neurons in the ventral and middle VTA also enriched medially (Figure 7D) and cluster 4 neurons in the middle and dorsal VTA confined to the most medial part of the VTA (Figure 7E). These findings indicate that VGluT2^+^ VGaT^+^, VGluT2^+^ VGaT^−^ and VGluT2^−^ VGaT^+^ neurons have a topographic organization more related to shared electrophysiological properties rather than to the neurotransmitter that they release.

**Figure 7.**
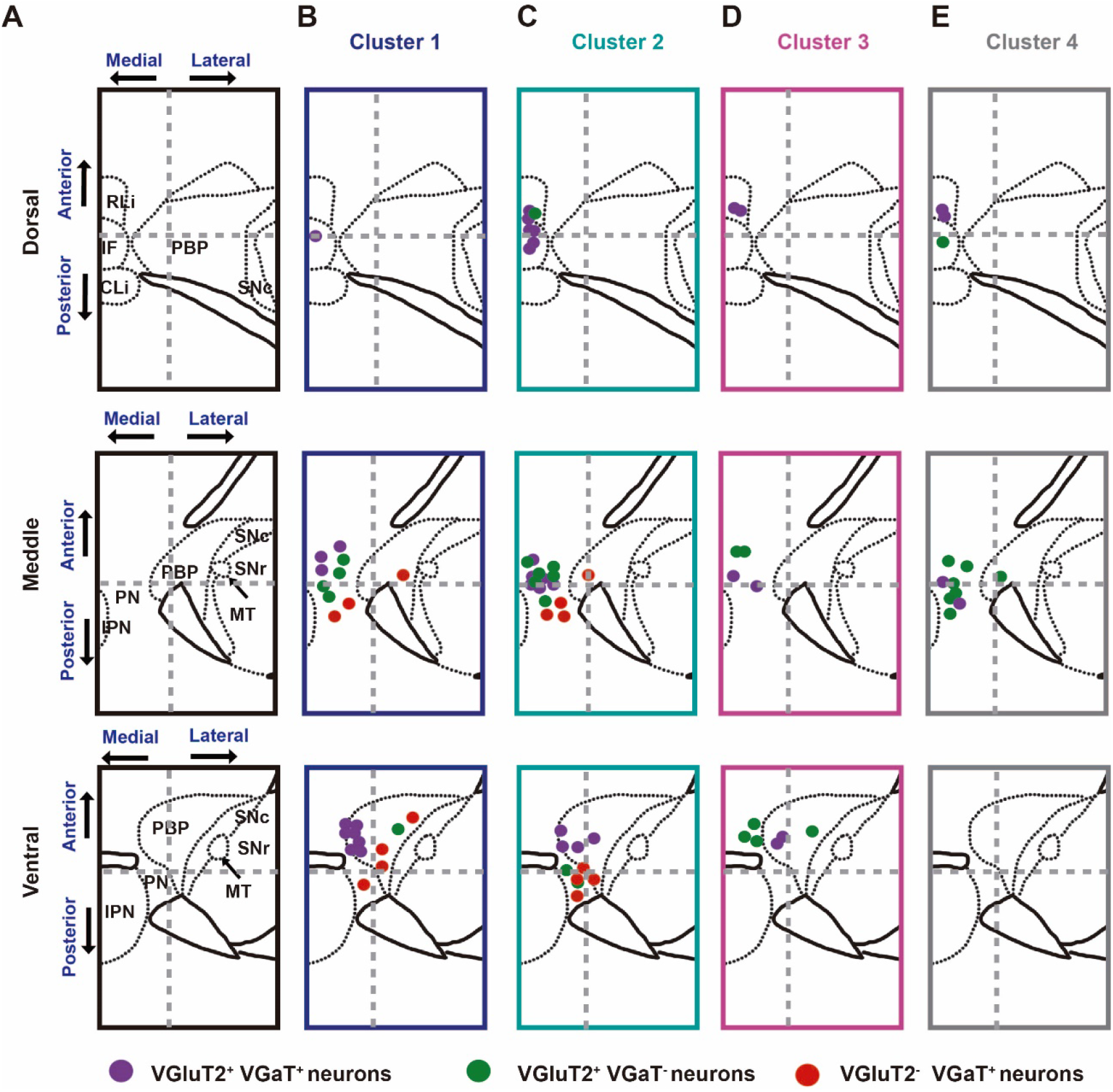
Topographic locations of VTA neurons by physiological cluster. **(A)** Schematic representations of dorsal, middle, and ventral horizontal VTA slices. (**B-E**) locations of recorded neurons from cluster 1 (**B**), cluster 2 (**C**), cluster 3 (**D**), and cluster 4 (**E**) in dorsal, middle, and ventral horizontal VTA slices. Grey dotted line divides the medial and lateral VTA. Each circle represents a single recorded neuron.

### μ-opioid receptors (MORs) are present in VTA GABA-releasing and glutamate-releasing neurons

Given that previous electrophysiological studies have demonstrated the presence of MORs in a subset of VTA GABA releasing neurons (Margolis et al., 2012), and anatomical studies have documented expression of Oprm1 mRNA in VTA VGluT2 neurons (Kudo et al., 2014), we systematically analyzed Oprm1 mRNA expression in VTA neurons that express VGluT2 mRNA and VGaT mRNA. We detected Oprm1 mRNA in VGluT2^+^ VGaT^−^ (Figure 8A), VGluT2^−^ VGaT^+^ (Figure 8B) and VGluT2^+^ VGaT^+^ neurons in coronal mouse sections (Figure 8C). We determined that within the total population of VTA neurons expressing Oprm1 mRNA (1,718 neurons, 3 mice), almost 20% expressed VGluT2 mRNA without VGaT mRNA (19.45% ± 0.9%; 336/1,718 neurons), 78% expressed VGaT mRNA without VGluT2 mRNA (78% ± 0.9%; 1,337/1,718 neurons), and a small number co-expressed VGluT2 mRNA and VGaT mRNA (2.5% ± 0.2%; 43/1,718 neurons) (Figure 8D). These findings demonstrated that GABA-releasing and glutamate-releasing neurons are two major classes of VTA neurons with the capability to synthesize MORs.

**Figure 8.**
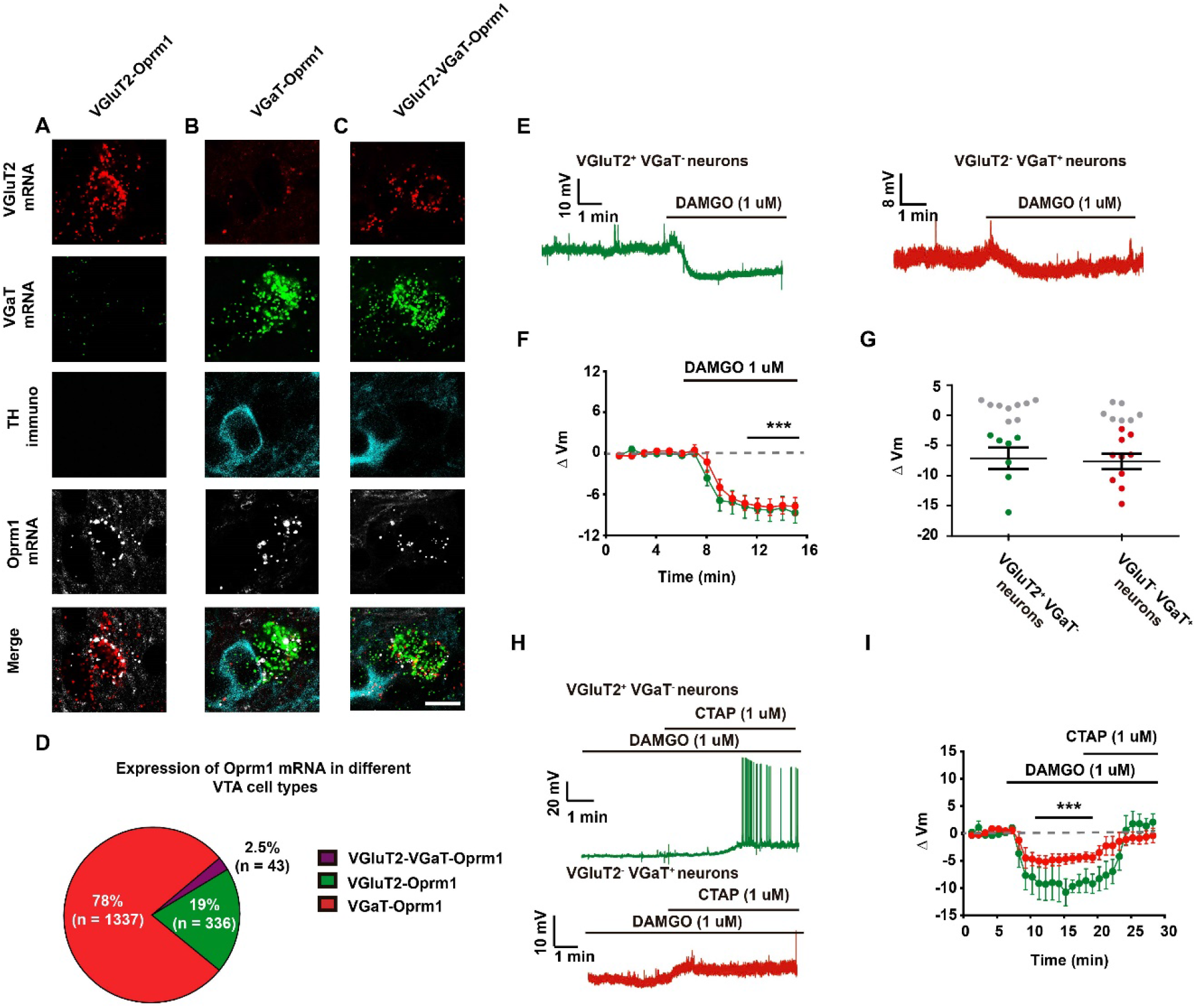
Functional MORs are present in VGluT2^+^ VGaT^−^ and VGluT2^−^VGaT^+^ VTA neurons. Detection of mRNA encoding VGluT2 (red), VGaT (green) or Oprm1 (white) and TH protein (cyan). **(A)** example neuron co-expressing VGluT2 and Oprm1 mRNAs. **(B)** example neuron co-expressing VGaT and Oprm1 mRNAs. **(C)** example neuron co-expressing VGluT2, VGaT, and Oprm1 mRNAs. **(D)** Percentage of VTA neurons expressing VGluT2 or VGaT mRNA with Oprm1 mRNA (3 mice). **(E)** Example current clamp traces from a VGluT2^+^ VGaT^−^(left) and a VGluT2^−^VGaT^+^ (right) neuron responsive to DAMGO bath application **(F)** Time course average showing membrane potential of VGluT2^+^ VGaT^−^ and VGluT2^−^VGaT^+^ neurons in response to DAMGO. **(G)** Summary of DAMGO-induced changes in membrane potential in VGluT2^+^ VGaT^−^ and VGluT2^−^VGaT^+^ neurons. Responses in green and red circles and nonrespondes in grey circles. Paired *t*-test before and after DAMGO application *t* 6= 4.042 p= 0.0068 for VGluT2^+^ VGaT^−^ neurons, *t*10= 5.991 p= 0.0002 for VGluT2^−^ VGaT^+^ neurons. **(H)** Example current clamp traces showing depolarizations in response to the μ-opioid receptor selective antagonist CTAP in neurons that were hyperpolarized by DAMGO. **(I)** Time course of average membrane potential in neurons with application of DAMGO and CTAP. Repeated measures ANOVA F2,14= 15.0 p˂0.0003, Dunnett’s multiple comparison test p˂ 0.0001 for VGluT2^+^ VGaT^−^; F2,17= 3.638 p˂0.0011, Dunnett’s multiple comparison test p˂ 0.0001 for VGluT2^−^VGaT^+^.

Next, we tested both VGluT2^+^ VGaT^−^ and VGluT2^−^ VGaT^+^ neurons for responses to the MOR selective agonist DAMGO (1μM). We detected DAMGO induced hyperpolarizations in a subset of VGluT2^+^ VGaT^−^ neurons (−7.1 ± 1.7 mV; −64.3 ± 2.7 mV baseline, −71.4 ± 3.2 mV DAMGO, n = 7 out of 17 tested neurons, 14 mice) (Figure 7 F-G). Hyperpolarizations were also observed in the presence of the GABA_A_ receptor antagonist (Bicuculline, 10μM) (Figure 8-supplement figure 1). Application of the MOR selective antagonist CTAP (1μM) reversed the DAMGO-induced hyperpolarizations (−73.3 ± 3.1 mV baseline, −84.5 ± 2.7 mV DAMGO, −72.5 ± 3.4 mV for DAMGO + CTAP, 5 tested neurons) (Figure 7 H-I). Similarly, we detected DAMGO induced hyperpolarizations in just over half of VGluT2^−^ VGaT^+^ tested neurons (−7.7 ± 1.3 mV; −64.1 ± 1.9 mV baseline, −71.8 ± 2.1 mV DAMGO, n =10 out of 18 tested neurons, 15 mice) (Figure 7 F-G), and these were also reversed by CTAP (−58.9 ± 1.0 baseline, −64.9 ± 1.2 DAMGO, −60.5 ± 1.2 DAMGO + CTAP, 6 tested neurons) (Figure 7 H-I). Collectively, these findings indicate that a subset of VTA neurons that release either GABA or glutamate express functional MOR in their somatodendritic region, the activation of which results in their hyperpolarization.

## Discussion

The VTA has historically been considered a dopamine brain structure, and the properties of these dopamine neurons have been intensively investigated for decades. However, the VTA has three additional major classes of neurons: GABA-releasing (VGluT2^−^VGaT^+^), glutamate-releasing (VGluT2^+^ VGaT^−^) and glutamate-GABA co-releasing (VGluT2^+^ VGaT^+^) neurons, whose physiological properties were unclear. To specifically study these three classes of neurons, we selectively tagged each class by *in vivo* expression of eYFP after intra-VTA delivery of INTRSECT viral vectors (C_ON_/F_ON_, C_ON_/F_OFF_ or C_OFF_/F_ON_) in different cohorts of double transgenic *vglut2-Cre/vgat-Flp* mice. We validated the selective expression of eYFP in each of the three targeted classes of VTA neurons by demonstrating that: (1) most of the VTA transduced neurons with C_ON_/F_ON_ viral vector (for targeting glutamate-GABA co-releasing neurons) co-expressed VGluT2 and VGaT mRNAs, (2) those transduced with C_ON_/F_OFF_ viral vector (for targeting glutamate-releasing neurons) expressed VGluT2 mRNA without VGaT mRNA, and (3) those transduced with C_OFF_/F_ON_ viral vector (for targeting GABA-releasing neurons) expressed VGaT mRNA without VGluT2 mRNA. Using *ex vivo* VTA recordings of the three classes of transfected neurons, we found that both glutamate-releasing and glutamate-GABA co-releasing neurons have lower excitability and lower basal firing activity than GABA-releasing neurons. In addition, while we observed diversity in the depolarization-induced firing patterns of glutamate-releasing and glutamate-GABA co-releasing neurons, the responses among GABA-releasing neurons were more uniform. We also demonstrated that whereas most of the VTA neurons containing the μ opioid receptors (MORs) are GABA-releasing neurons, 40% of glutamate-releasing neurons were also hyperpolarized by MOR activation. Collectively, we provide evidence that: (1) the VTA neuronal firing of both glutamate-releasing and glutamate-GABA co-releasing neurons require a stronger excitatory input to fire than GABA-releasing neurons, (2) the ionic channel composition is likely to be more diverse among glutamate-releasing and glutamate-GABA co-releasing neurons than among GABA-releasing neurons, and (3) postsynaptic MOR activation inhibits the activity of both GABA-releasing and glutamate-releasing VTA neurons.

It is well documented that *I*_h_ is present in both dopamine and non-dopamine neurons (Jones and Kauer, 1999; Margolis et al., 2006), including neurons expressing GAD (Chieng et al., 2011; Margolis et al., 2012; Ntamati et al., 2018), VGaT (Woodward et al., 2019) or VGluT2 (Hnasko et al., 2012). We extended these observations by showing that *I*_h_ is present in less than half of the glutamate-GABA co-releasing neurons, about half of the glutamate-releasing neurons and in more than 90% of the GABA-releasing neurons. However, the meane *I*_h_ magnitude is smallest in glutamate-GABA co-releasing neurons, followed by GABA-releasing neurons, and largest in glutamate-releasing neurons. We found that regardless of the neuronal cell type, the neurons with the greatest *I*_h_ magnitudes were located in the lateral VTA, and those with low amplitudes were in the medial VTA. These findings are consistent with the VTA latero-medial neuronal heterogeneity observed in dopamine neurons (Li X et al.,2013, Morales and Margolis 2017) and show it is a property shared by all classes of VTA neurons. While the molecular bases underlying differences in *I*_h_ magnitude among VTA neurons remains to be determined, one possibility is differential levels of expression of any of the four hyperpolarization-activated cyclic nucleotide-gated channel (HCN1-4) subunits that generate *I*_h_ and whose transcripts have been detected in the VTA (Monteggia et al., 2000). As an alternative, differences in *I*_h_ magnitude may reflect differential neuronal distribution of the HCN channels across neuronal compartments, as the HCN subunits have been detected in the plasma membrane of cell bodies, dendrites, or axons (Notomi and Shigemoto, 2004).

While our findings on the intrinsic properties of VTA neurons suggest that stronger excitatory drive is required for firing in VTA glutamate-releasing and glutamate-GABA co-releasing neurons compared to VTA GABA-releasing neurons, it remains to be determined which glutamatergic sources target each class of VTA neuron. Ultrastructural and electrophysiological reports indicate that VTA neurons expressing either GAD or VGaT receive excitatory input from different brain areas. For instance, pioneer ultrastructural studies showed that VTA GABA-neurons (expressing GAD) receive asymmetric (excitatory-type) synapses from axon terminals whose neurons originate in the lateral habenula (Olmelchenko et al., 2009), medial prefrontal cortex (Carr and Sesack, 2000), periaqueductal grey (Olmelchenko and Sesack, 2010), and bed nucleus of the stria terminalis (Kudo et al., 2012). Furthermore, recent findings utilizing optogenetics and VTA slice electrophysiology have shown that the firing of GABA neurons (expressing VGaT or GAD) is evoked by exciting glutamatergic inputs (expressing VGluT2) from lateral hypothalamus neurons (Nieh et al., 2015), superior colliculus neurons (Zhou et al., 2019) or periaqueductal grey neurons (Ntamati et al., 2018). Moreover, a circuitry-based study on input from the periaqueductal grey to VTA showed that periaqueductal glutamatergic neurons preferentially target the GAD neurons with large *I*_h_ (Ntamati et al., 2018). Though it is possible that some of the GABA neurons identified in these prior studies were in fact glutamate-GABA co-releasing neurons, information on glutamatergic afferents to VTA glutamatergic neurons is limited. We recently reported quantitative ultrastructural, optogenetic, and electrophysiological evidence that VTA glutamate-releasing neurons (VGluT2^+^ VGaT^−^) receive a strong glutamatergic input from the lateral hypothalamic area, LHA (Barbano et al., 2020). Furthermore, we also showed that the somatodendritic region of a single VTA glutamate-releasing neuron receives multiple asymmetric synapses from axon terminals arising from LHA glutamatergic neurons (Barbano et al., 2020). Thus, these observations underscore the importance that characterization of the precise synaptic organization between different classes of VTA neuronal types and specific afferents, from neurons of the same nature and shared brain structure, provides on achieving a better understanding on the firing regulation of diverse classes of VTA neurons.

Previous studies found hyperpolarization-induced rebound burst firing in subsets of VTA dopamine and non-dopamine neurons mediated by *I*_h_ (Tateno and Robinson, 2011) or T-type calcium channels (Tracy et al., 2018, Woodward et al., 2019). We found the same types of responses in subpopulations of glutamate-GABA co-releasing, glutamate-releasing and GABA-releasing neurons. A recent VTA electrophysiological study in a VGaT Knock-in rat line expressing the fluorescent protein Venus reported two populations with different rebound firing properties (type 1 with low threshold calcium spikes during rebound firing, and type two with post hyperpolarizing action potentials during rebound firing) of VGaT-Venus neurons expressing T-channels (Woodward et al., 2019). Given that we detected T-channel mediated rebound in glutamate-GABA co-releasing and GABA-releasing neurons, and both classes of neurons express VGaT (Root et al., 2018b), it remains to be determined the extent to which these two classes of neurons overlapped with those detected in rat VTA VGaT-Venus neurons.

Findings from electrophysiological and pharmacological studies show presynaptic and postsynaptic MOR function in the VTA (Johnson and North 1992, Margolis et al., 2012; Fields and Margolis, 2015). While postsynaptic MORs are generally thought to be limited to GABA neurons within the VTA, we observed transcripts encoding MORs (Oprm1) expressed in a subset of glutamate-releasing neurons, and these neurons were clustered in the midline aspects of the VTA. We also found that MOR activation hyperpolarizes these glutamate-releasing neurons. These findings of postsynaptic actions of MORs within two subpopulations of VTA neurons together with the presence of MORs in synaptic terminals (Margolis et al., 2004, Zhang et al., 2015, Chen et al., 2015, Bull et al., 2017) underscore the complex actions of opioids within the VTA.

In summary, we detected unique as well as overlapping electrophysiological properties among the glutamate-GABA co-releasing, glutamate-releasing and GABA-releasing VTA neurons. Our electrophysiological findings indicate that firing of VTA glutamate-GABA co-releasing and glutamate-releasing neurons may require stronger excitatory drive compared to the GABA-releasing neurons. However, given that the neuronal firing pattern depends on both the intrinsic properties of the neurons and the network activity innervating them, future studies are necessary to identify the origin, nature, and impact of inputs to the specific classes of VTA neurons.

## Materials and Methods

### Experimental subjects

Both male and female mice were used in this study. The *vglut2-IRES::Cre* mice (JAX # 016963) and *vgat::FlpO* mice (JAX # 031331; Daigle et al., 2018) were crossed to produce a *vglut2-IRES::Cre x vgat::FlpO* mice. Animals were housed in temperature- and humidity-controlled facilities under a 12 h light/dark cycle with dawn at 0700 h and ad libitum chow and water prior to the start of experimental procedures. Mice were 2-3 months of age at the start of the experiment. Experiments were conducted in accordance with the USPHP Guide for the Care and Use of Laboratory Animals and approved by the Animal Care and Use Committee of the National Institute on Drug Abuse Intramural Research Program.

### Surgery and Virus Injections

Mice were anesthetized with isoflurane (2-4% induction; 1% maintenance) and secured to a stereotaxic frame. After exposing the top of the skull, the mouse’s head was leveled to ensure the skull was flat. One of the following 3 viruses were injected into the VTA (0.3 μl; AP: −3.1 to −3.3, ML: ± 0.0, DV: −4.3 to −4.4) to label the different classes of VTA neurons: (1) AAV5-Hsyn-C_ON_-F_ON_-eYFP to label VGluT2^+^ VGaT^+^ neurons, (2) AAV5-Hsyn-C_ON_-F_OFF_-eYFP to label VGluT2^+^ VGaT^−^ neurons or (3) AAV5-Hsyn-C_OFF_-F_ON_-eYFP to label VGluT2^−^ VGaT^+^ neurons. Injections were made using a Micro4 controller and UltraMicroPump along with 10 μl Nanofil syringes equipped with 35-gauge needles (WPI Inc., Sarasota, FL). Syringes were left in place for 10 min following injections to minimize diffusion. Following surgery, mice recovered on a warm heating pad before being transferred back to the vivarium home cage. Mice remained in the colony to allow for recovery and virus expression for 6-8 weeks for RNAscope experiments and 4-6 weeks prior electrophysiology experiments.

### Combination of RNAscope *in situ* Hybridization and Immunolabeling

Tissue preparation: *Wild-type* mice and *Vglut2-IRES::Cre x Vgat::FlpO* mice of six to eight weeks following virus injections were anesthetized with chloral hydrate (0.5 ml/kg) and perfused transcardially with 4% (w/v) paraformaldehyde (PF) in 0.1 M phosphate buffer (PB), pH 7.3. Brains were left in 4% PF for 2 h and transferred to 18% sucrose in PB overnight at 4°C to prepare them for RNAscope *in situ* hybridization-immunohistochemistry experiments. The detection of transcripts encoding VGluT2 mRNA, VGaT mRNA and Oprm1 mRNA were done by using RNAscope, and TH detection by immunohistochemistry. Coronal free-floating sections (wild-type mouse, VTA, 16 μm) were incubated for 2 h at 30°C with Mouse anti-TH antibody (1:1000, MAB318, Millipore, Burlington, MA) in DEPC-treated phosphate buffer (PB) with 0.5% Triton X-100 supplemented with RNasin (Promega, Madison, WI). Sections were rinsed 3 × 10 min with DEPC-treated PB, and incubated with secondary Donkey anti-Mouse Alexa Fluor 750 (1:100, ab175738, abcam, Cambridge, MA) for 1 h at 30°C. Sections were rinsed with DEPC-treated PB and then were mounted onto Fisher SuperFrost slides and dried overnight at 60°C. RNAscope *in situ* hybridization processing was performed in accordance to the manufacturer’s instructions (Advanced Cell Diagnostics, Newark, CA). Briefly, sections were treated with heat and protease digestion followed by hybridization with a mixture containing target probes to mouse VGluT2 (319171), VGaT (319191-C3) and Oprm1 (489311-C2). Additional sections were hybridized with the bacterial gene DapB as a negative control, which did not exhibit fluorescent labeling. VGluT2 were detected by Atto 550, VGaT were detected by Alexa 488, and Oprm1 were detected by Atto 647. GFP immunolabeling and detection of mRNA for VGluT2 and VGaT were performed as described above. VTA Coronal free-floating sections (*VGluT2-Cre x VGaT-FlpO* mouse, 16 μm in thickness) were processed for immunodetection of Mouse anti-GFP antibody (1:500, 632381, Takara Bio USA, Inc. Mountain View, CA) and incubated with secondary Donkey anti-Mouse Alexa Fluor 488 (1:100, 715-545-151, Jackson ImmunoResearch, West Grove, PA), after processing by RNAscope *in situ* hybridization, VGluT2 were detected by Atto 550, VGaT were detected by Atto 647. RNAscope *in situ* hybridization sections were viewed, analyzed, and photographed with an Olympus FV1000 confocal microscope or a Keyence BZ-X800 microscope. Negative control hybridizations showed negligible fluorophore expression. Neurons were counted when the stained cell was at least 5 μm in diameter. Pictures were adjusted to match contrast and brightness by using Adobe Photoshop (Adobe Systems). The number of mice (n=3/group; 13-16 sections/mouse) analyzed was based on previous studies in our lab using radioactive detection of VGluT2 mRNA from rat VTA neurons.

### Patch-clamp recordings

Six to eight weeks after virus injection, mice were anesthetized with isoflurane, decapitated, and the brain was quickly removed and placed in ice-cold artificial cerebrospinal fluid (ACSF), saturated with 95% O_2_ and 5% CO_2_, and modified to contain (in mM): 92 NMDG, 20 HEPES, 25 glucose, 30 NaHCO_3_, 1.2 NaH_2_PO_4_, 2.5 KCl, 5 sodium ascorbate, 3 sodium pyruvate, 2 thiourea, 10 MgSO_4_, 0.5 CaCl_2_ on a VT-1200 vibratome (Leica, Nussloch, Germany), and sectioned through the VTA in horizontal slices (200 μm thick). The slices were placed in a holding chamber filled with the same solution but held at 32°C. After 10-15 minutes, slices were transferred to a holding chamber containing room temperature ACSF modified to contain (in mM): 92 NaCl, 20 HEPES, 25 glucose, 30 NaHCO_3_, 1.2 NaH_2_PO_4_, 2.5 KCl, 5 sodium ascorbate, 3 sodium pyruvate, 2 thiourea, 1 MgSO_4_, 2 CaCl_2_. For recordings, slices were transferred to a chamber superfused with 32°C ACSF containing (in mM): 125 NaCl, 2.5 KCl, 1.25 NaH_2_PO_4_, 1 MgCl_2_, 2.4 CaCl_2_, 26 NaHCO_3_, and 11 glucose. Electrodes (2-4MΩ) were backfilled with an internal solution containing (in mM): 120 potassium gluconate, 8.0 NaCl, 1.0 MgCl_2_, 10 HEPES, 2.0 Mg-ATP, 0.3 Na_2_-GTP, 10 ditris-phosphocreatine, 0.2 EGTA and 0.2% biocytin (pH 7.2; 275-290 mOsm). Cells were visualized on an upright microscope using infrared differential interference contrast video microscopy. Whole-cell voltage clamped, and current clamped recordings were made using a MultiClamp 700B amplifier (2 kHz low-pass Bessel filter and 10 kHz digitization) with pClamp 10.3 software (Molecular Devices, Sunnyvale, CA). Firing rate was determined before breaking into the cell under cell attached mode for at least 60 seconds of continuous activity. *I*_h_ was measured under voltage clamp mode holding at −60 mV and stepping to −120, −100, and −80 mV for 1000 ms. Membrane potential and AP properties were measured in current clamp I=0 within the first 5 minutes after gaining whole cell access. To determine excitability and firing properties, an input/output curve consisting of depolarizing current steps of 500 ms duration from 0-150 pA were applied and the number of APs fired during each current step were quantified with pClamp 10.1 software. To determine adaptation of AP firing, ISI of the first 10 AP fired after current injection able to evoke the maximum response was analyzed. Neurons with an increase of more than 50% of the ISI during the first 10 AP was classified as neuron with adaptation. *I*_A_ current was measured in voltage clamp using two step protocol. The first step consisted of a 500 ms hyperpolarizing pre-pulse (−120 mV, 500 ms) followed by increasing depolarizing steps from −100 mV to 30 mV for 1000 ms. The second step was a depolarizing pre-pulse (40 mV) followed by depolarizing steps from −100 mV to 30 mv 1000 ms. *I*_A_ was obtained by subtracting the currents generated by the second protocol from the currents generated by the first protocol. In the current clamp recordings where *I*_A_ was blocked by 4-AP, the membrane potential of the cell was clamped in the membrane potential before the addition of 4-AP and CNQX was applied to prevent an increase AP firing due to an increase in glutamatergic transmission.

### Cluster analysis

Cluster analysis was applied to data from 131 VTA neurons using 21 electrophysiological parameters that include: membrane potential, membrane capacitance, membrane resistance, time constant, tonic firing frequency, Coefficient of variation of tonic firing frequency, Ih current amplitude, rheobase, action potential threshold, gap between resting membrane potential and action potential threshold, AP amplitude, action potential duration, after hyperpolarized potential amplitude, after hyperpolarized potential duration, after hyperpolarized potential peak, latency to fire action potential at rheobase stimulation, number of action potentials fired at rheobase, number of action potentials fired after hyperpolarization, maximum number of action potentials fired after depolarization, current to induce maximum number of action potentials fired, highest firing frequency. Data from each electrophysiological parameter was normalized to its mean value to prevent over representation of a specific parameter during clustering. Cluster analysis was performed in R-studio software using K-means method and Euclidean distance. The result of the clustering was plotted as dendrogram and a hierarchical tree. A principal component analysis was applied to determine the electrophysiological parameters that best separate the clusters and comparisons between clusters were made using one-way ANOVA and Tukey’s *posthoc* test.

### Statistical analysis

Data in the figures are presented as mean ± SEM, one-way ANOVA or student’s t-test were used to compare group of neurons using prism 5.0 software. P< 0.05 was required for significance.

## Acknowledgments

This work was supported by the Intramural Research Program of the National Institute on Drug Abuse. We thank Drs. David Root and Francois Vautier for setting the colony of *Cre/Flp* transgenic mice at NIDA/IRP. We also thank Dr. Nirnath Sah (JHU) for advice on cluster analysis

## Author Contributions

MM and JM-B. conceptualized and initiated the project. JM-B and IC performed electrophysiological studies. JM-B, IC, GEM-S and EM analyzed electrophysiological data. SM, HW, BL performed immunolabeling studies and quantified neurons from RNAscope studies. H-LW and BL performed RNAscope studies. SZ performed confocal studies and data analysis. MM and JM-B and EM prepared the manuscript with contribution from all authors.

---

**Figure 4-figure supplement 1.**
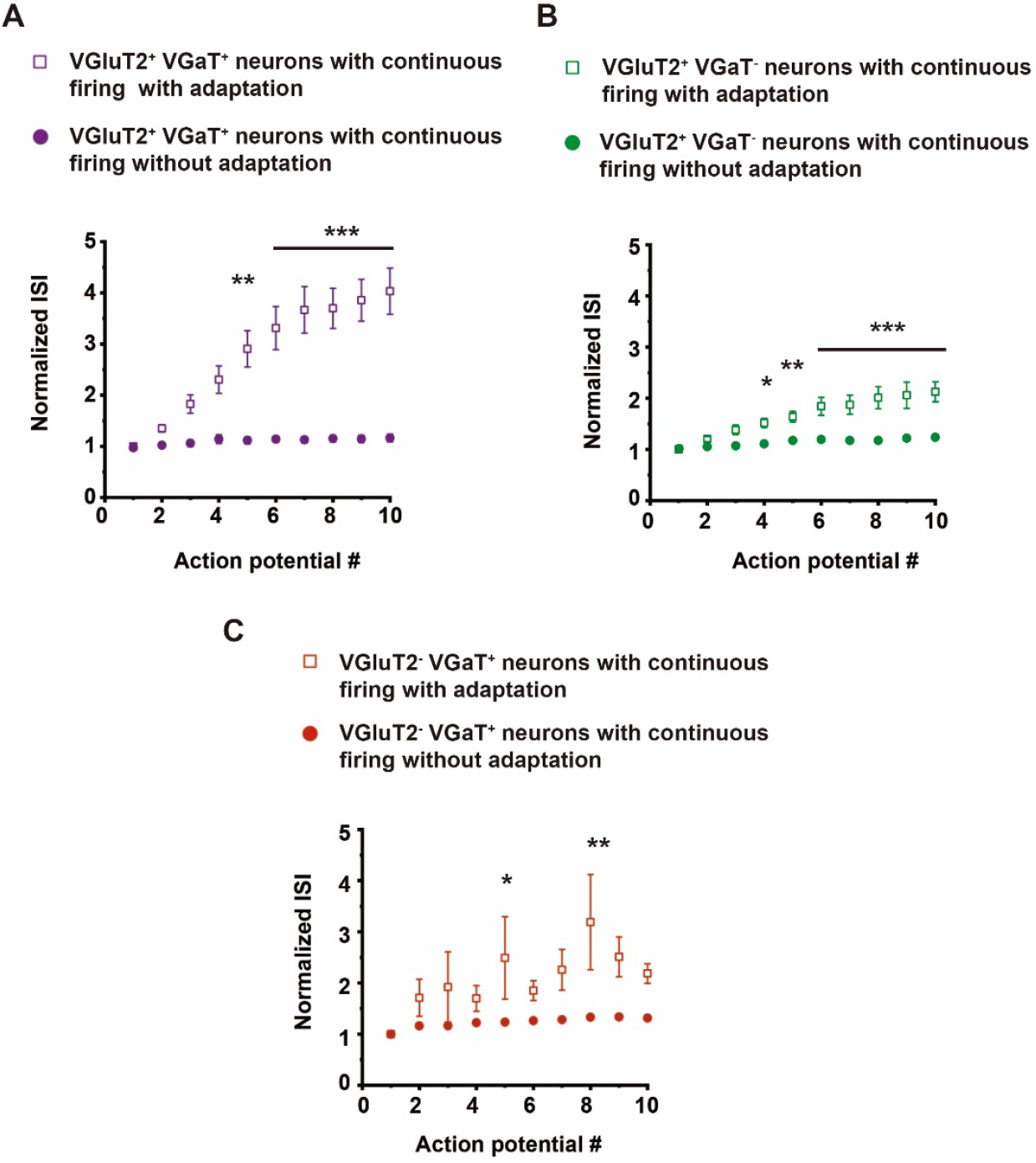
Neurons that fire with adaptation during depolarizing current steps have larger inter-spike intervals (ISIs) than neurons without adaptation. **(A-C)** Normalized ISI of the first 10 AP fired after 150 pA current injection from **(A)** VGluT2^+^ VGaT^+^ neurons with adaptation and without adaptation. Two way ANOVA spike number x type of neuorn F_9,250_= 4.72 p˂ 0.0001, type of neuron F_1,250_= 121.44 p˂ 0.0001, spike number F_9,250_= 5.73 p˂ 0.0001 Bonferroni posthoc test **p˂ 0.01, ***p˂ 0.001. **(B)** VGluT2^+^ VGaT-neurons with adaptation and without adaptation Two way ANOVA spike number x type of neuorn F_9,220_= 5.08 p˂ 0.0001, type of neuron F_1,220_= 147.51 p˂ 0.0001, spike number F_9,220_= 11.31 p˂ 0.001. Bonferroni posthoc test *p= 0.05, **p˂ 0.01, ***p˂ 0.001 and **(C)** VGluT2^−^VGaT^+^ neurons with adaptation and without adaptation Two way ANOVA spike number x type of neuorn F_9,360_= 1.39 p= 0.1904, type of neuron F_1,360_= 38.80 p˂ 0.0001, spike number F_9,360_= 2.48 p˂ 0.094. Bonferroni posthoc test *p= 0.05, **p˂ 0.01.

**Figure 4-figure supplement 2.**
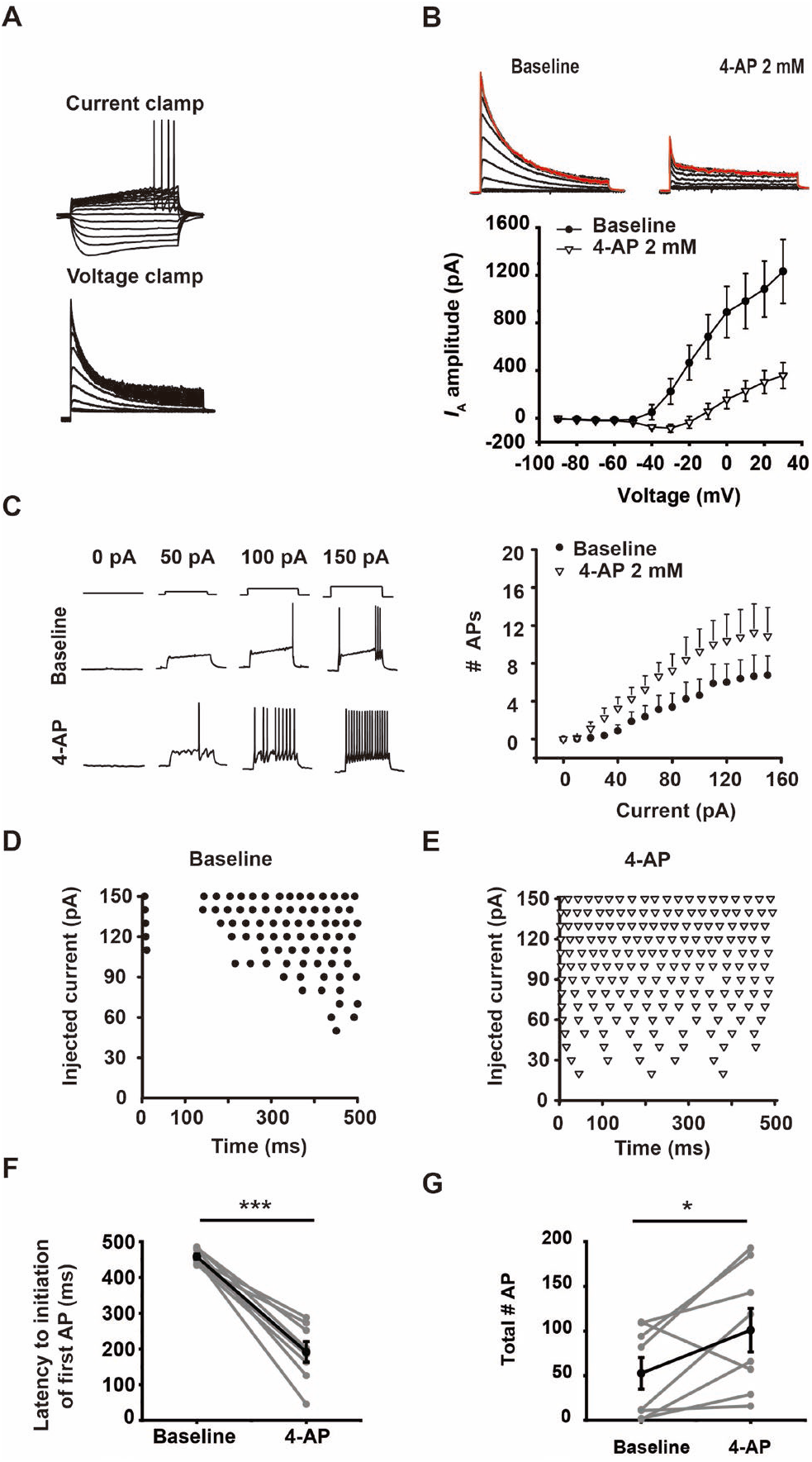
*I*_A_ mediates long latency AP firing. **(A)** Example **c**urrent clamp (top) and voltage clamp (bottom) traces in a VGluT2^+^ VGaT-long latency neuron **(B)** Example voltage clamp traces from a VGluT2^+^ VGaT^−^ long latency neuron at baseline and after bath application of the *I*A blocker 4- Aminopyridine (4-AP) (top). Mean *I* A amplitude responses across different voltage steps at baseline (close circles) and after bath application of 4-AP (open triangles) (bottom) (n= 12 neurons, 3 VGluT2^+^ VGaT^+^ neurons and 9 VGluT2^+^ VGaT-) **(C)** Example current clamp traces (left) of a long latency VGluT2^+^ VGaT^+^ neuron during depolarizing step current injections before and after bath application of 4-AP. Input/output curves show 4-AP (open triangles) increased number of APs fired during depolarizing current injections compared to baseline (closed circles) (n= 8 neurons, 4 VGluT2^+^ VGaT^+^ neurons and 4 VGluT2^+^ VGaT-neurons) Two way ANOVA 4-AP x current F_15,112_= 0.83 p= 0.6408, 4-AP F_15,,112_= 49.06 p˂ 0.0001, current F_15,112_= 4.64 p˂ 0.0001. **(D-E)** Example firing pattern during injected current steps in a VGluT2^+^ VGaT^+^ long latency neuron at baseline **(D)** and after bath application of 4-AP (E). **(F)** 4-AP decreased the latency to AP firing during the minimum injected current step in long latency neurons (n= 8 neurons, 4 VGluT2^+^ VGaT^+^ and 4 VGluT2^+^ VGaT^−^). Paired *t*-test *t*7= 8.877 ***p˂ 0.0001 **(G)** 4-AP increased total number of APs fired after activation across long latency neurons during an input/output curve (10 – 150 pA, 500 ms). Paired *t*-test *t*7= 2.426 *p= 0.0457 (n= 8 neurons 4 VGluT2^+^ VGaT^+^ and 4 VGluT2^+^ VGaT-)

**Figure 5-figure supplement 1.**
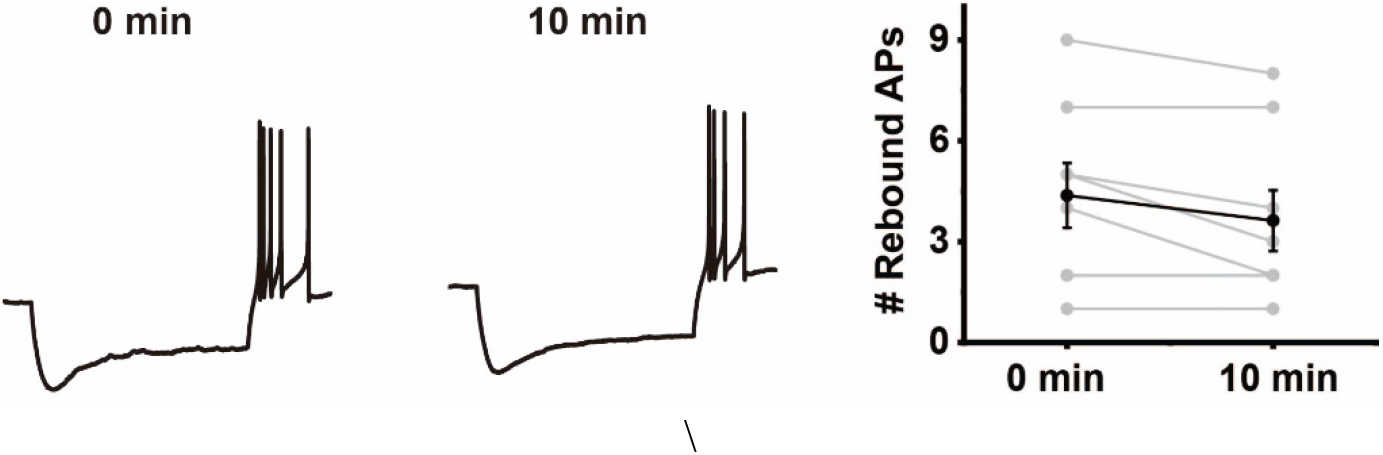
Rebound firing is stable across time. **(A)** Current clamp traces from a VGluT2^−^VGaT^+^ neuron with rebound firing at the beginning of the experiment (0 minutes) and after 10 minutes. **(B)** Number of rebound APs in at the beginning of the experiment (0 minutes) and after 10 minutes of recording (n= 8 neurons, 2 VGluT2^+^ VGaT^+^ neurons, 2 VGluT2^+^ VGaT-neurons, and 4 VGluT2^−^VGaT^+^ neurons)

**Figure 8-figure supplement 1.**
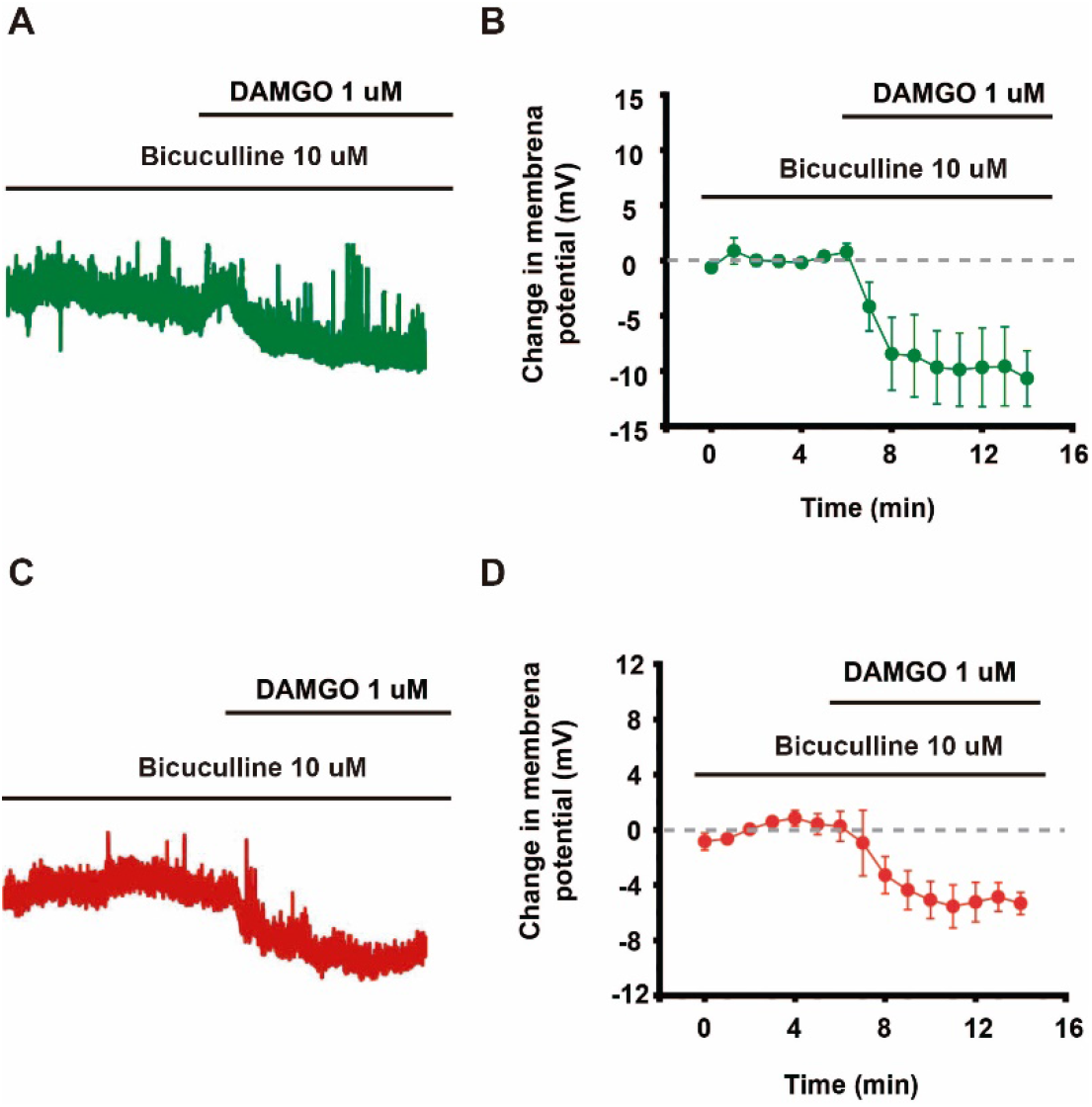
VGluT2^+^ VGaT^+^ and VGluT2^−^VGaT^+^ neurons are inhibited by DAMGO in the presence of GABA_A_ receptor antagonist Bicuculline. **(A)** Example current clamp of a VGluT2^+^ VGaT-neuron that responded to DAMGO in the presence of GABAA receptor antagonist Bicuculline. **(B)** Mean time course showing the change in membrane potential across VGluT2^+^ VGaT-neurons (n= 4) to application of DAMGO in the presence of Bicuculline. **(C)** Example current clamp trace of a VGluT2^−^VGaT^+^ neuron that responded to DAMGO in the presence of the GABAA receptor antagonist Bicuculline. **(D)** Mean time course of the membrane potential of VGluT2^−^VGaT^+^ neurons in response to DAMGO in the presence of Bicuculline (n= 5).

